# Imaging active site chemistry and protonation states: NMR crystallography of the tryptophan synthase α-aminoacrylate intermediate

**DOI:** 10.1101/2021.05.12.443852

**Authors:** Jacob B. Holmes, Viktoriia Liu, Bethany G. Caulkins, Eduardo Hilario, Rittik K. Ghosh, Victoria N. Drago, Robert P. Young, Jennifer Romero, Adam D. Gill, Paul M. Bogie, Joana Paulino, Xiaoling Wang, Gwladys Riviere, Yuliana K. Bosken, Jochem Struppe, Alia Hassan, Jevgeni Guidoulianov, Barbara Perrone, Frederic Mentink-Vigier, Chia-en A. Chang, Joanna R. Long, Richard J. Hooley, Timothy C. Mueser, Michael F. Dunn, Leonard J. Mueller

**Author notes:** co-first authors (Jacob B. Holmes, Viktoriia Liu, Bethany G. Caulkins). Corresponding authors: Leonard J. Mueller, Michael F. Dunn. **Author Contributions:**. Designed research: LJM, MFD. Performed research: JBH, VL, BGC, EH, RKG, VND, RPY, JR, JP, XW, GR, YKB, JS, AH, JG, BP, FMV. Contributed new reagents/analytic tools: ADG, PMB. Analyzed data: LJM, MFD, TCM, RJH, JRL, CAC. Wrote paper: LJM, MFD, JBH, VL, BGC, RKG. The authors declare that they have no known competing financial interests.

## Abstract

NMR-assisted crystallography – the synergistic combination of solid-state NMR, X-ray crystallography, and first-principles computational chemistry – holds remarkable promise for mechanistic enzymology: by providing atomic-resolution characterization of stable intermediates in the enzyme active site – including hydrogen atom locations and tautomeric equilibria – it offers insight into structure, dynamics, and function. Here, we make use of this combined approach to characterize the α-aminoacrylate intermediate in tryptophan synthase, a defining species for pyridoxal-5′-phosphate-dependent enzymes on the β-elimination and replacement pathway. By uniquely identifying the protonation states of ionizable sites on the cofactor, substrates, and catalytic side chains, as well as the location and orientation of structural waters in the active site, a remarkably clear picture of structure and reactivity emerges. Most incredibly, this intermediate appears to be mere tenths of angstroms away from the preceding transition state in which the β-hydroxyl of the serine substrate is lost. The position and orientation of the structural water immediately adjacent to the substrate β-carbon suggests not only the fate of the hydroxyl group, but also the pathway back to the transition state and the identity of the active site acid-base catalytic residue. Reaction of this intermediate with benzimidazole (BZI), an isostere of the natural substrate, indole, shows BZI bound in the active site and poised for, but unable to initiate, the subsequent bond formation step. When modeled into the BZI position, indole is positioned with C3 in contact with the α-aminoacrylate C^β^ and aligned for nucleophilic attack.

**Significance Statement:** The determination of active site protonation states is critical to gaining a full mechanistic understanding of enzymatic transformations; yet hydrogen positions are challenging to extract using the standard tools of structural biology. Here we make use of a joint solid-state NMR, X-ray crystallography, and first-principles computational approach that unlocks the investigation of enzyme catalytic mechanism at this fine level of chemical detail. For tryptophan synthase, this allows us to peer along the reaction coordinates into and out of the α-aminoacrylate intermediate. Through this process, we are developing a high-resolution probe for structural biology that is keenly sensitive to proton positions – rivaling that of neutron diffraction, yet able to be applied under conditions of active catalysis to microcrystalline and non-crystalline materials.

## Introduction

Pyridoxal-5′-phosphate (PLP), Scheme 1, participates in numerous enzyme catalyzed reactions essential for amino acid metabolism, including transamination, decarboxylation, and α/β/γ-elimination and substitution (5–8). The power of PLP as a cofactor comes from its ability to act as an electron sink, allowing for the stabilization of carbanionic intermediates. A more subtle aspect of PLP catalysis demonstrated in β-elimination and replacement reactions is the ability of the cofactor to fine tune the polarity at the β-carbon of amino acids, facilitating the elimination of poor leaving groups and their replacement with weak nucleophiles.

**Scheme 1:**
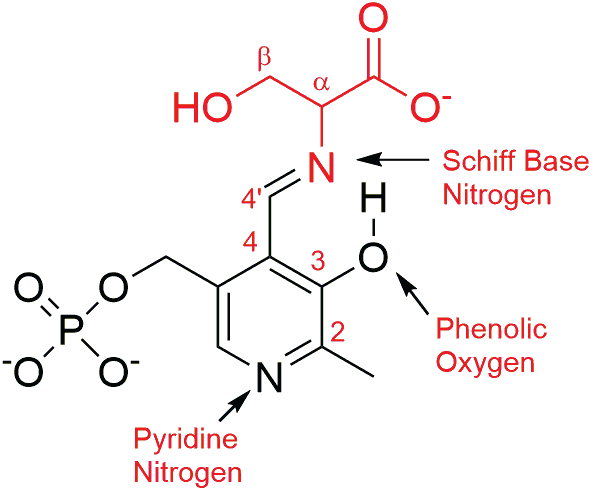
External aldimine intermediate with PLP in black and L-Ser substrate in red

Tryptophan synthase is the prototypical example of a PLP-dependent enzyme on the β-elimination and replacement pathway. The *Salmonella typhimurium* tryptophan synthase *(StTS)* studied here is a 143 kDa αββα bienzyme complex (9). TS catalyzes the last two steps in the biosynthesis of L-Tryptophan: the α-subunit cleaves indole-3-glycerol 3’-phosphate (IGP) to glyceraldehyde-3-phosphate (G3P) and indole (the α-reaction), while the β-subunit catalyzes the PLP-dependent β-elimination and replacement of the L-Ser hydroxyl group with indole to produce L-Trp (the β-reaction, Fig. 1) (10, 11). The fidelity of the proton transfers in the TS catalytic cycle are critical for maintaining the β-elimination and substitution pathway. It has been proposed that this specificity has its origins in the modulation of protonation states and tautomeric equilibria of the PLP cofactor, which are in turn directed by chemical interactions with acid-base groups within the active site (12, 13). Hence, the catalytic protein residues interacting with the cofactor establish the appropriate chemical and electrostatic environment to favor a particular protonation state and reaction pathway (6, 8, 13).

**Fig. 1.**
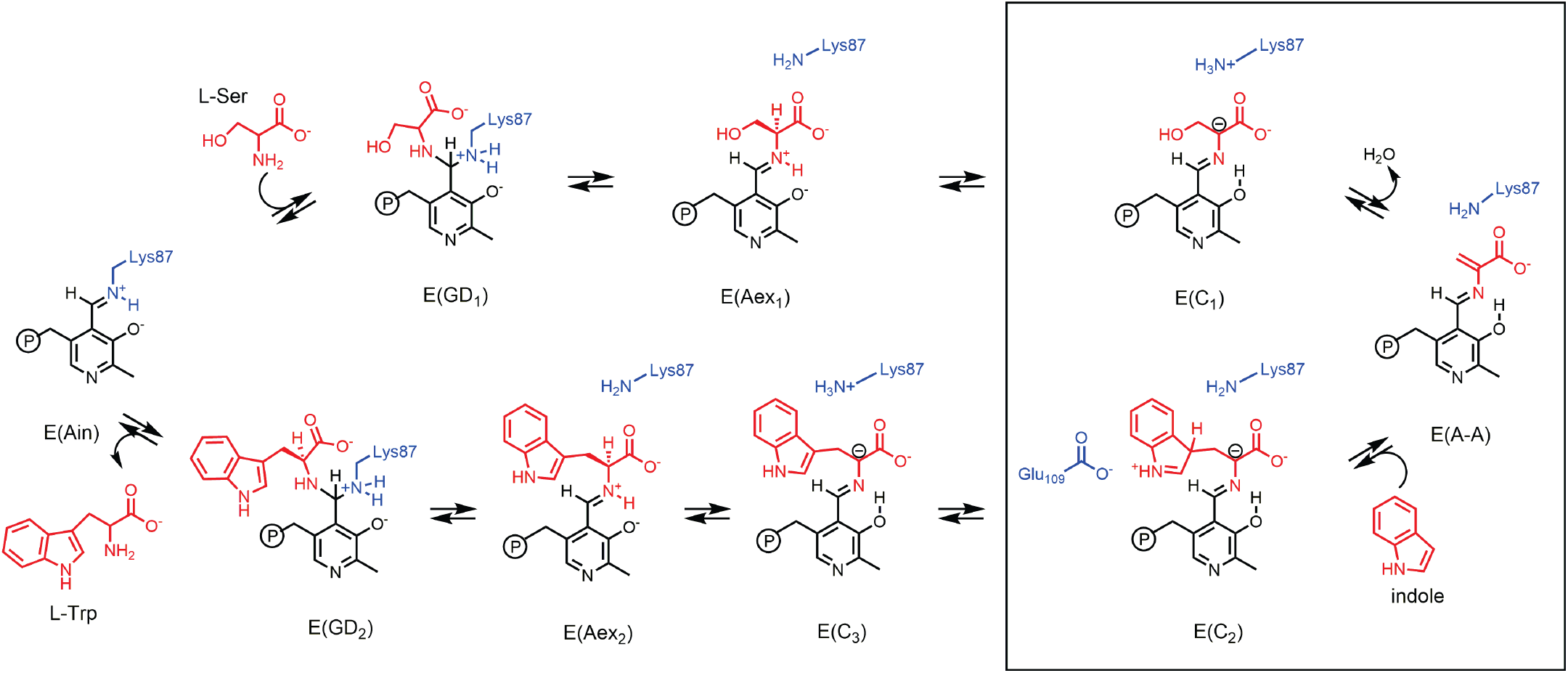
Tryptophan Synthase β-Site Reaction, highlighting the α-aminoacrylate intermediate (1, 2). In stage I of the reaction (top), L-Ser reacts with the internal aldimine, E(Ain), to give the gem-diamine, E(GD1), and external aldimine, E(Aex_1_), intermediates. Subsequent proton abstraction via βLys87 leads to the first carbanionic intermediate, E(C_1_), and elimination of the β-hydroxyl group as water gives the α-aminoacrylate intermediate, E(A-A). In stage II (bottom), indole makes a nucleophilic attack at C^β^ of E(A-A) to yield a new C-C bond and the L-Trp carbanionic intermediate, E(C_2_), and, upon deprotonation, E(C_3_). In the final stages, E(C_3_) is reprotonated, leading to the eventual release of L-Trp and the regeneration of E(Ain).

The determination of protonation states of enzyme intermediates, however, remains a significant challenge to the tools of structural biology. Even high-resolution X-ray crystal structures are challenged to place hydrogen atoms. Recently, there have been tremendous strides in neutron crystallography, which in joint refinement with X-ray crystallography can define both heavy atom and hydrogen atom locations (14, 15). But the drawback to neutron crystallography is the requirement for large, preferably perdeuterated, crystals and multi-week-long acquisition times at room temperature. The latter currently precludes analysis of reactive intermediates such as the ones studied here, although next generation high intensity neutron sources promise to ameliorate this challenge. Cryo-EM is also pushing to the boundaries of resolution necessary for hydrogen atom detection (16), a remarkable achievement, yet currently far from routine.

In joint application with NMR spectroscopy, cryo-EM and diffraction methods continue to push towards complete, atomic-resolution descriptions of structure and function (17–20). For delineating the chemistry of the active site, NMR and diffraction become even more powerful when combined with first-principles computational chemistry. We are developing NMR-assisted crystallography – the joint SSNMR, X-ray, and first-principles computational refinement of crystal structures – to solve for the chemically-rich three-dimensional structures of enzyme active sites (21–23). By chemically-rich we mean structures where the location of all atoms, including hydrogen, are specified. NMR crystallography was originally developed within the context of molecular organic and inorganic solids (24–28). More recently, our group (21–23) and others (29–31) have been working to extend this approach to structural biology, where it can provide consistent and testable models of structure and function of enzyme active sites. Our approach is three-fold: X-ray crystallography is used to provide a coarse structural framework upon which chemically-detailed models of the active site are built using computational chemistry, and various active site chemistries explored; these models can be quantitatively distinguished by comparing their predicted NMR chemical shifts with the results from solid-state NMR experiments. Provided a sufficient number of chemical shift restraints are measured within the active site, NMR-assisted crystallography can uniquely identify a consistent structure or, equally important, determine that none of the candidates is consistent with the experimental observations. For the case of the “quinonoid” intermediates in tryptophan synthase, this approach demonstrated that the intermediate is actually better described as a carbanionic species with a deprotonated pyridine ring nitrogen – a structure that we propose is fundamental to understanding reaction specificity in TS (23).

Here, we make use of NMR-assisted crystallography to characterize the TS α-aminoacrylate intermediate, a species that marks a major divergent step in PLP chemistry as only enzymes that perform β-elimination reactions generate this intermediate (32, 33). In this case, NMR-assisted crystallography is able to uniquely identify the protonation states of ionizable sites on the cofactor, substrates, and catalytic side chains, as well as locate and orient structural waters in the active site. From this, a remarkably detailed picture of structure and reactivity emerges in which the α-aminoacrylate intermediate appears to be tenths of angstroms away from the preceding transition state in which the serine β-hydroxyl is lost. The position and orientation of the structural water immediately adjacent to the substrate suggests the fate of the β-carbon hydroxyl group, as well as the pathway back to the transition state and the identity of the active site acid-base catalytic residue. Subsequent characterization of the Michaelis complex of the α-aminoacrylate intermediate with the indole isostere, benzimidazole (BZI), shows BZI bound in the active site and poised for, but unable to initiate, the successive bond formation step. When modeled into the BZI position, the indole C-3 makes contact with C^β^ of the α-aminoacrylate moiety and is aligned for nucleophilic attack at the C-C bond formation step, while the amino group of βLys87 is correctly positioned for transfer of the proton from the indole C-3 carbon to the C^α^ carbanion in the next steps. Here, the chemically-detailed, three-dimensional structure from NMR-assisted crystallography is key to understanding why BZI does not react, while indole does.

## Results and Discussion

### X-ray Crystallography

X-ray crystal structures for the TS aminoacrylate intermediate, E(A-A), and the TS TS aminoacrylate-BZI complex, E(A-A)(BZI), have been reported previously by our group (34) and others (35). Formation of the E(A-A) intermediate is characterized by adoption of closed conformations by both the α-subunit and the β-subunit (1, 11, 34–37), triggered by the reaction of L-Ser with the PLP cofactor to produce the E(A-A). This stimulates the α-site reaction by at least 30-fold, as the β-subunit active site becomes poised to fuse indole to the PLP-substrate complex (38).

Fig. 2A shows the β-subunit active site from the crystal structure of the E(A-A) intermediate. Three crystallographic waters are seen adjacent to the serine substrate, forming a hydrogen bonded chain extending from the carboxylate of the catalytically essential βGlu109 (39). A similar set of waters was also observed in E(A-A) crystal structures 4HN4 (35) and our new crystal structure 7MT4 (vide infra). The position of the central water is particularly striking, as it forms close contacts to both the substrate C^β^ (C-O distance of 3.2 Å) and the βLys87 ε-amino group (N-O distance of 3.0 Å). This places it in close proximity to the site that the β-hydroxyl group of E(C1) is expected to occupy so that βLys87 can hydrogen bond with, and transfer a proton to, the hydroxyl oxygen in the preceding mechanistic step. The binding of BZI (Fig. 2B) displaces the waters and causes a small perturbation of the α-aminoacryloyl group of E(A-A). The binding also induces small movements of the βGlu109 side chain carboxyl and the βLys87 ε-amino group so that they are within hydrogen bonding distances of the BZI N1 and N3, respectively. These interactions and the close structural similarity of BZI to indole make the BZI complex a good mimic of the expected alignment of indole for nucleophilic attack on E(A-A) C^β^. Absent in Fig. 2, however, are the hydrogen atoms, which are critical to delineating the hydrogen bond donors and acceptors and their mechanistic roles.

**Fig. 2.**
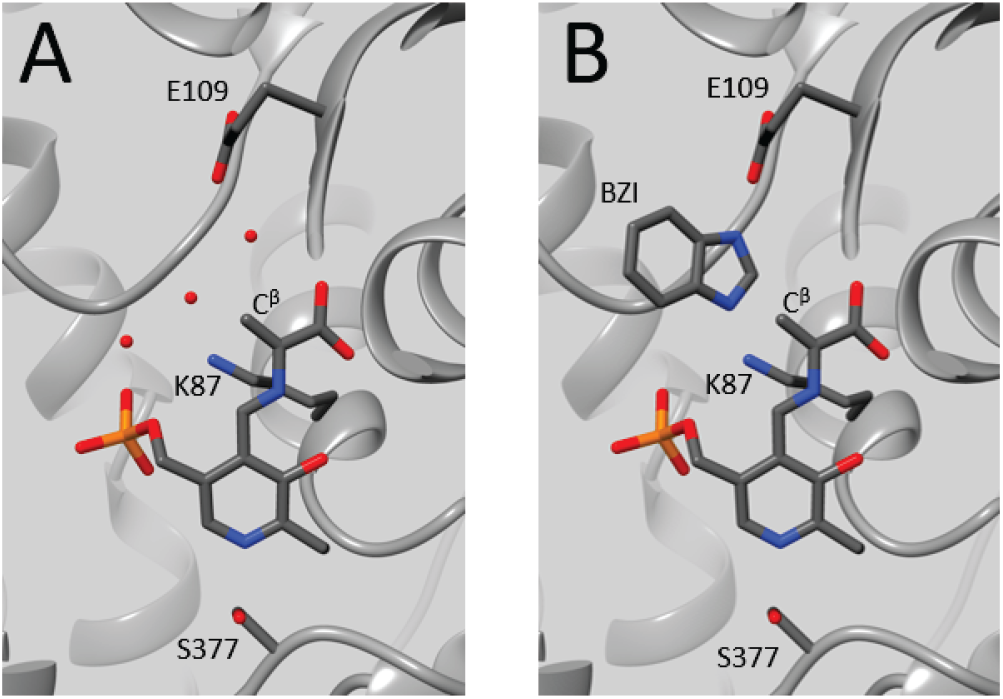
The β-subunit active sites for E(A-A) and E(A-A)(BZI) from crystal structures 2J9X and 4HPX, respectively. (A) The E(A-A) intermediate shows 3 active site waters adjacent to the substrate with the central water forming close contacts to both the substrate C^β^ and the βLys87 ε-amino group. (B) The E(A-A)(BZI) complex shows BZI displacing the three waters, but otherwise inducing only small changes in the active site structure. Images rendered in UCSF Chimera (4).

### Solid-State NMR

Comprehensive understanding of the mechanism in TS requires the full chemical structure of each intermediate in the catalytic pathway. To address this, we turn to solid-state NMR: the chemical shift in NMR is an extremely sensitive reporter on protonation and hybridization states, and SSNMR can provide chemical shifts under closely analogous conditions to those for the acquisition of the diffraction data. Both the E(A-A) and E(A-A)(BZI) complexes can be observed by SSNMR under conditions that favor the accumulation of the enzyme-bound species while the microcrystals remain catalytically active (40, 41). The α-aminoacrylate intermediate can be established by supplying only the L-Ser substrate, or L-Ser and the inhibitor BZI. However, a competing side reaction eventually converts L-Ser to ammonium and pyruvate. Solid-state NMR spectra of the reaction of L-Ser with microcrystalline TS to form the α-aminoacrylate intermediate are shown in Fig. 3. The CPMAS spectra correspond to solid-state crystalline protein and bound substrate. The acquisition of these spectra was interleaved with single pulse, low-power decoupling experiments reporting predominantly on free ligand and reaction products in the surrounding mother liquor (Fig. S4, SI). The α-aminoacrylate intermediate persists as long as there is excess serine in solution, after which the corresponding resonances in the solid-state NMR spectra decay.

**Fig. 3.**
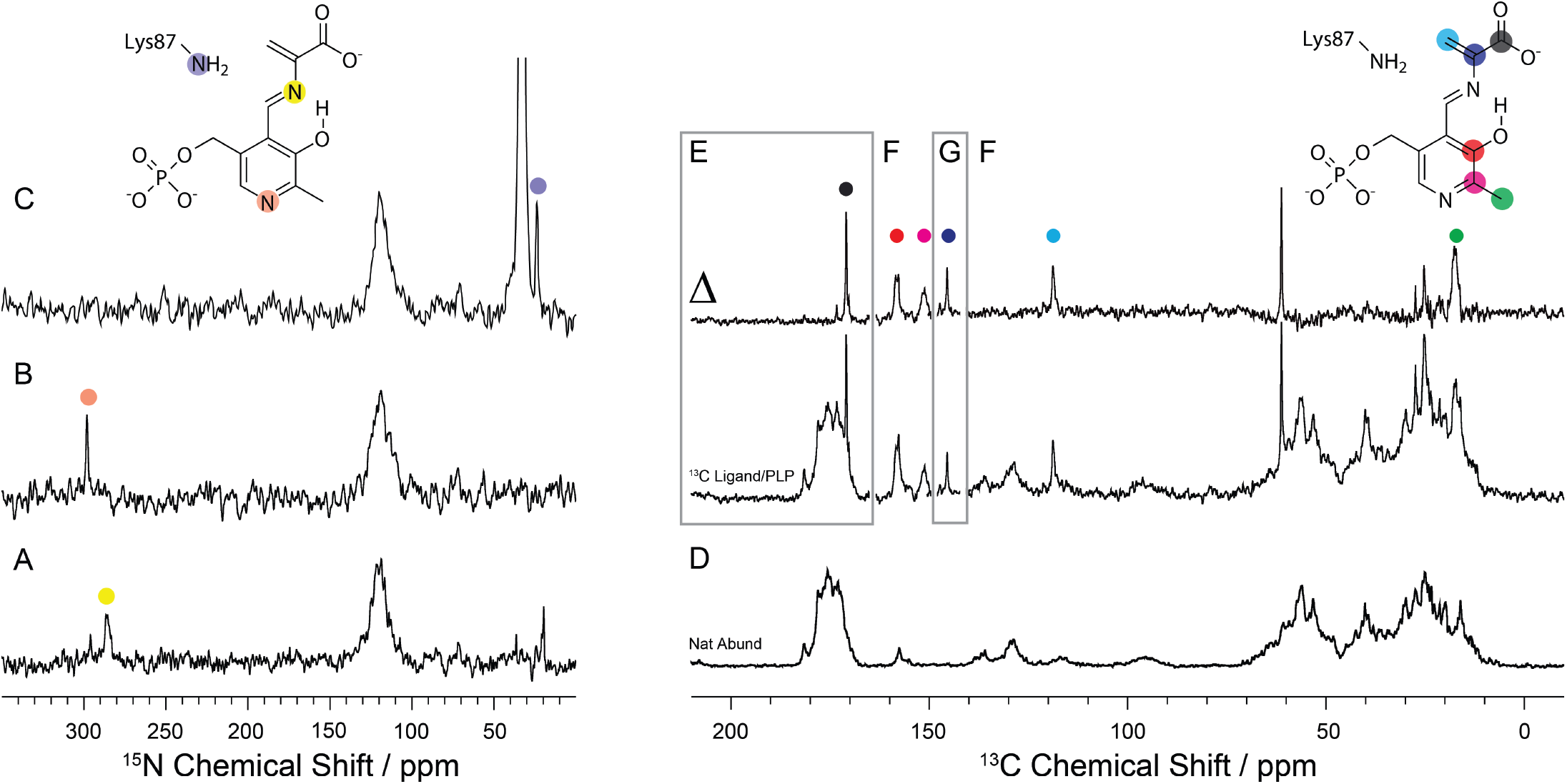
^15^N and ^13^C SSNMR CPMAS spectra of microcrystalline TS E(A-A) prepared with the following isotopic labeling: (A) ^15^N-labeled on the substrate L-Ser; (B) selectively ^15^N enriched on the PLP cofactor; (C) selectively ^15^N-enriched at protein lysine side chain ε-amino groups; (D) natural abundance isotopomer concentration; (E) ^13^C-labeled on C’ of the L-Ser substrate; (F) selectively ^13^C,^15^N-enriched on the PLP cofactor and C^β^ of the substrate L-Ser; and (G) ^13^C-labeled on C^α^ of the substrate L-Ser. The top spectra in (E)-(G) are formed as the difference between the E(A-A) spectra with various cofactor/ligand isotopic labels and the same spectra acquired at natural abundance, highlighting the resonances for the specific site labels. The large peak at 63.1 ppm is free serine. Spectra acquired at 9.4 T, −10 °C, and 8 kHz MAS as described in the Methods and Materials.

The solid-state NMR spectra for E(A-A) shown in Fig. 3 were acquired with various combinations of ^15^N and ^13^C labels on the PLP cofactor (Fig. 3B,F), L-Ser substrate (Fig. 3A,E,F,G), and the protein lysine ε-amino nitrogen sites (Fig. 3C). With unlabeled/natural abundance substrate and protein (Fig. 3D), the spectra show only unresolved background signals from the protein, including the ^15^N signal from the backbone amides (centered ~120 ppm in 2A-C) and ^13^C resonances for the backbone carbonyl (~165-175 ppm), aliphatic (~10-70 ppm), and aromatic (~110-160 ppm) carbons. With the “magnification of the isotopic enrichment” (24) for the various components, distinct resonances for the active site species become visible and can be uniquely assigned.

The spectra of the sample prepared with ^15^N and ^13^C on the PLP cofactor (Fig. 3B,F) show a nitrogen resonance at 297.6 ppm and three additional carbon resonances at 17.5, 151.2, and 158.1 ppm that are assigned to the cofactor N1, C2’, C2, and C3, respectively. The sample prepared using ^13^C and ^15^N-L-Ser as a substrate (Fig. 3A,E,F,G) displays a nitrogen resonance at 286.7 ppm and additional carbon resonances at 170.9 ppm, 145.6 ppm, and 118.8 ppm that are assigned to the Schiff base nitrogen and carbons that derive from the serine C’, C^α^, and C^β^, respectively. The Schiff base nitrogen was found to have a significant temperature dependence of ~-0.14 ppm/K, while the PLP ring nitrogen showed no discernable temperature dependence (Fig. S5). The full chemical shift tensor was measured for the Schiff base nitrogen under slow MAS conditions as shown in Fig. S6; the principal axis components, {δ_11_, δ_22_, δ_33_} = {526.5, 306.3, 31.5} ppm, were extracted by fitting the intensity of the spinning sideband manifold (42).

These shifts help establish several key elements of the E(A-A) intermediate’s chemical structure. At this point in the catalytic cycle, the serine substrate has lost its β-hydroxyl and there is a double bond between C^α^ and C^β^, which is confirmed by their chemical shifts of 145.6 ppm and 118.8 ppm, respectively. Second, at 286.7 ppm, the chemical shift of the Schiff base nitrogen indicates a neutral imine linkage, which is further supported by the large span of the chemical shift tensor (43). This isotropic chemical shift, however, is below the limiting value for a fully neutral Schiff base, and the substantial temperature dependence (−0.14 ppm/K) suggests tautomeric exchange involving transient protonation of this site (13, 23). At the same time, the PLP C2 and C3 shifts of 151.2 and 158.1 ppm indicate that the PLP phenolic oxygen is protonated (neutral) (23). Finally, the PLP nitrogen shift of 297.6 ppm is indicative of a neutral (deprotonated) pyridine ring nitrogen (N1) for the cofactor. The shift is somewhat upfield compared to model compounds in solution (12), but stands in stark contrast to the corresponding shift in the Fold Type I PLP-dependent aspartate aminotransferase, in which N1 falls at 213 ppm (adjusted to the δ_N_(liq-NH_3_) scale), indicating that it is predominantly protonated (13). This change in protonation at N1 correlates with the PLP ring hydrogen bonding partner switching from aspartic acid to serine when moving from AAT to TS.

The ^15^N chemical shift for the catalytic lysine side chain βLys87 was measured by the preparation of a TS sample in which all the protein lysine residues were ^15^N-enriched at the ε-amino sites (ε-^15^NH_3_-Lys TS) as reported previously (44). In the initial resting state, E(Ain), this sample shows a peak at 202 ppm, corresponding to the protonated Schiff base linkage between βLys87 and the cofactor, and a large number of overlapped resonances centered at 33 ppm, assigned to charged lysine ε-amino groups that are predominantly solvent-exposed at the surface of the protein. Upon the addition of L-Ser to the microcrystalline sample, the resonance at 202 ppm is lost and a new resonance at 24.2 ppm is observed (Fig. 3C). Based on this time correlation, this resonance is assigned to the ε-amino group of active site βLys87. This chemical shift suggests a neutral amino group for βLys87 consistent with its proposed role as the primary acid-base catalytic residue: at this point in the catalytic cycle (Fig. 1), βLys87 has delivered a proton to the β-hydroxyl of the serine substate, which is eliminated as water, and is now neutral and poised to act as a base and abstract a proton once indole adds to form E(C_2_).

The ^31^P isotropic chemical shift and chemical shift tensor were measured under CPMAS conditions for the cofactor’s phosphate group (Fig. S7). The isotropic chemical shift and the skew of the shift tensor show a characteristic response in going from the mono-to the di-anionic charge state. The phosphate isotropic shift of 5.2 ppm and its full shift tensor point to a dianionic charge state, in keeping with other TS intermediates (23, 44) and many other PLP dependent enzymes (45).

Finally, the ^17^O chemical shifts of the substrate carboxylate group were measured using ^17^O quadrupole central transition (QCT) NMR in solution (46, 47) (Fig. S2); these were the only shifts measured for the intermediate in solution, all others were measured in the solid state for microcrystalline sample preparations. ^17^O NMR is not yet considered a standard high-resolution probe for biomolecular NMR, but QCT NMR takes advantage of the unique property of the NMR central transition to narrow as the size of the protein-substrate complex increases. We had previously reported preliminary isotropic ^17^O shifts for both the E(A-A) and E(A-A)(BZI) complexes (46); here we take advantage of additional measurements on the 35.2 T series connect hybrid magnet at the National High Magnetic Field laboratory to improve the accuracy of the extracted parameters as described in the SI (48). The ^17^O chemical shifts of 257 ppm and 289 ppm are consistent with an ionized carboxylate group. The difference between the two shifts for E(A-A) reflects differences in the number and strength of their hydrogen bonding interactions with the residues in the carboxylate binding pocket (residues β110-114), with the upfield shifted signal tentatively assigned to the oxygen strongly hydrogen bonded to the side chain βThr110, which forms the closest contact.

Chemical shifts for the E(A-A)(BZI) complex were measured analogously (SI, Figs. S2-S9) and are summarized along with the E(A-A) shifts in Table 1. The chemical shifts for the complex formed in the presence of both serine and BZI are nearly identical to those for the intermediate formed with serine alone, with one prominent exception: the loss of the neutral amine resonance at 24.1 ppm and the appearance of an additional charged amino group resonance at 35.6 ppm. The latter was assigned to the ε-amino group of active site βLys87 based on ^15^N{^31^P}-rotational-echo double resonance (REDOR) (49) experiments that place it in close spatial proximity (<4 Å) to the phosphorus atom of the cofactor (Fig. S9). There are also two additional nitrogen resonances for the BZI inhibitor itself at 165.5 and 227.8 ppm that are assigned to the protonated and deprotonated ring nitrogen atoms, N1 and N3 (Table 1). Note that the shift in the charge and protonation state of βLys87 upon addition of the BZI inhibitor puts it out of sync with its mechanistic role as a base at this point in the catalytic cycle.

**Table 1.**
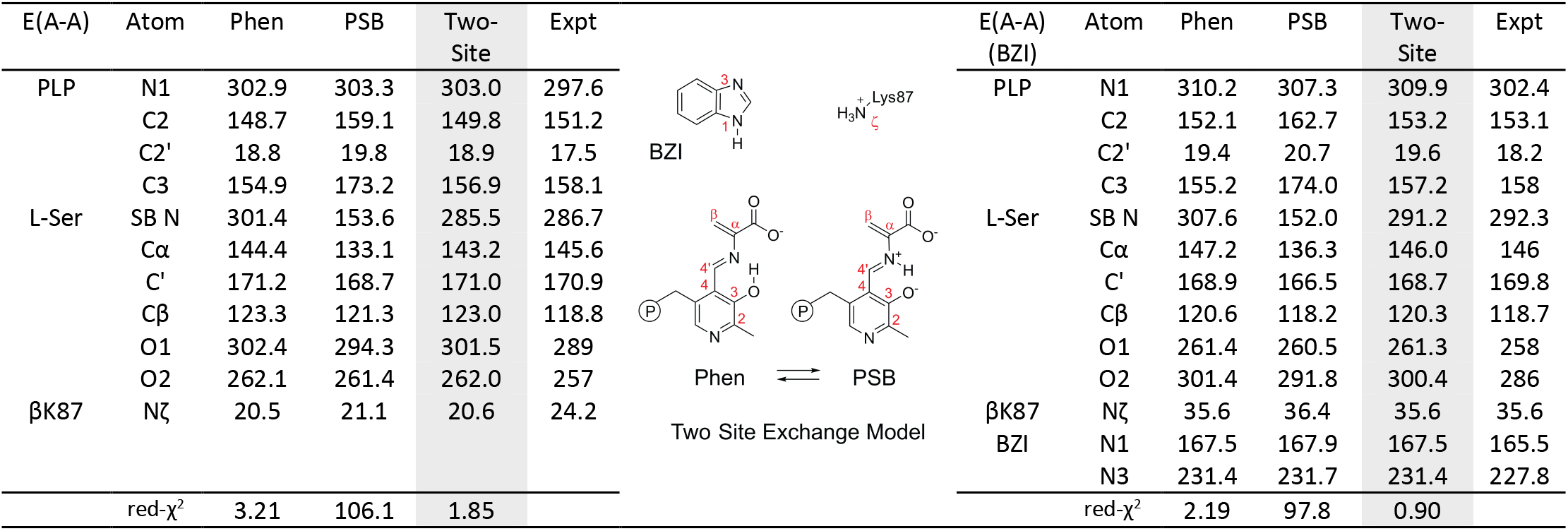
E(A-A) and E(A-A)(BZI) experimental and first-principles chemical shifts (ppm) for the phenolic (phen) and protonated Schiff base (PSB) models and their two-site exchange model with the following populations: E(AA) 89.3% Phen,10.7% PSB; E(AA)(BZI) 89.4% Phen, 10.6% PSB

### First-Principles Calculations: NMR-Assisted Crystallography

While the preliminary chemical structures above shed significant light on the chemistry of the active site, they are incomplete in two important respects: first they are absent detailed three-dimensional structure; second, the chemical shifts do not fully conform to the expected limiting shifts from model compound studies. To place the chemistry of the active site in fuller structural context, it is necessary to move beyond the empirical correlation of shift with environment, to the more quantitative methods of first-principles computational chemistry. This approach melds the chemical structure from NMR with the three-dimensional structural framework from crystallography, giving a chemically-detailed crystal structure of the enzyme active site in which the spatial coordinates of all atoms, including hydrogens, are specified. This process also allows for the quantitative testing of the proposed chemical structures, a full description of potential tautomeric exchange dynamics, and the determination of whether the description is unique and fully consistent with the experimental data.

The state-of-the-art for first-principles chemical shift calculations is now quite advanced, and extensive benchmarking by our group (50, 51) and others (52) has shown that if the correct structure is known – down to the position of each hydrogen atom and the orientation of each water and hydroxyl – then the NMR chemical shifts can be predicted to better than 1.5 ppm RMSD for carbon, 4.3 ppm for nitrogen, and 7.5 ppm for oxygen. This expected quantitative agreement allows us to establish a screening protocol in which proposed structures are evaluated (and ranked) for consistency with the experimental chemical shifts using the reduced chi-squared statistic (Eq. 4, Materials and Methods) to determine whether the proposed model has computationally predicted shifts that agree with the experimental chemical shifts to within the expected RMSD error ranges. We take as the benchmark for a solved structure the unique identification of a single structure (or a fast-exchange equilibrium between closely related tautomers) that satisfies the 95% confidence limits of the reduced-χ^2^. In practice, this requires multiple (e.g., 10-12) chemical shift measurements throughout the active site and supplemental chemical shift tensor and chemical shift temperature dependence coefficient measurements to uniquely identify a single model (23).

To introduce minimal bias into the structural analysis, an extensive pool of candidate structures was generated by systematically varying the protonation states of the ionizable sites on the cofactor, substrates, and catalytic βLys87 ε-amino group as shown in Fig. 4 and described in detail in the Materials and Methods (SI). The candidate structures were constructed directly as three-dimensional cluster models of the active site, built on the framework of the X-ray crystal structure, and included all residues and crystal waters within 7 Å of the substrate and cofactor. The atoms at the exterior of the cluster were pinned at their crystallographic locations and a fully quantum-mechanical, DFT-based geometry optimization was undertaken for the interior residues, substrate, and cofactor, and for the prediction of the NMR chemical shifts. The clusters contained ~703 atoms (Fig. 4B), a size for which the convergence and accuracy of the chemical shift calculations has been established (50).

**Fig. 4.**
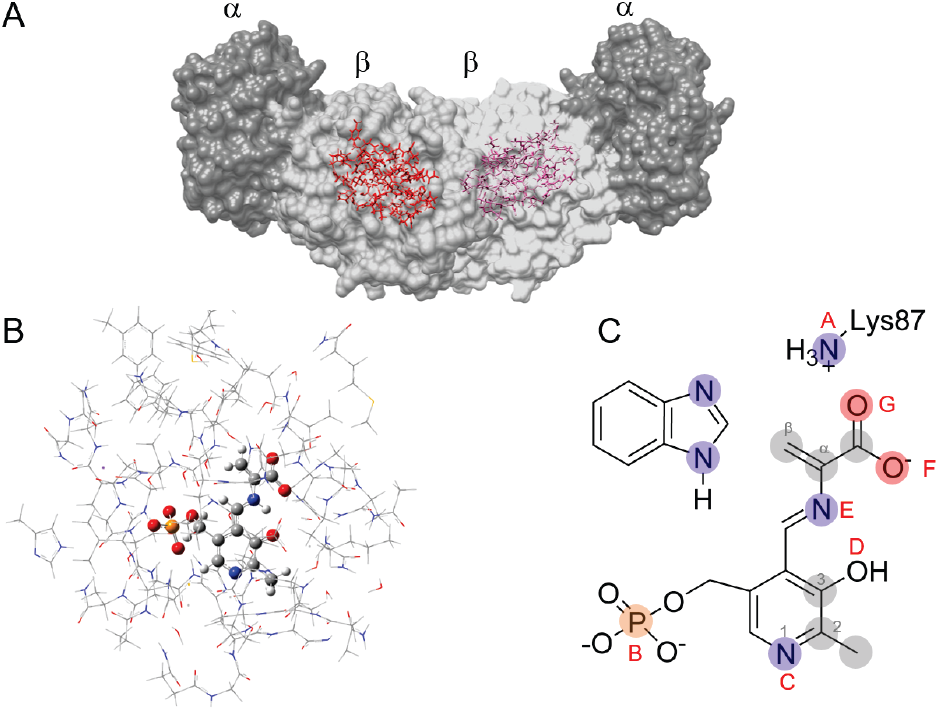
Cluster model of the E(A-A) active site. (A) X-ray crystal structure of the TS α2β2 heterodimer with the β-subunit active site in red. (B) Cluster model of the active site for first-principles geometry optimization and chemical shift calculations; protein side chains displayed in wireframe and cofactor and substrate in ball-and-stick. (C) Protonation sites on and near the cofactor/substrate complex: A, the βLys87 side chain; B. the PLP phosphate group; C, the PLP pyridine ring nitrogen; D, the PLP phenolic oxygen; E, the Schiff-base nitrogen; and F/G, the substrate carboxylate. Shaded nuclei indicate sites for which experimental NMR chemical shifts are reported.

35 initial models of E(A-A) (Scheme S1) and E(A-A)(BZI) (Scheme S2) were generated, geometry optimized, and their chemical shifts predicted, as summarized in Tables S1 and S2. The structures were then ranked based on their agreement with the experimental chemical shifts as quantified by the reduced-χ^2^ statistic as shown in Fig. 5. There is a nice differentiation of models in both cases, yet the reduced-χ^2^ values for both fall outside the expected value for 95% confidence, indicating that these single static structures of the active site are incomplete and do not reproduce the expected chemical shifts within the accuracy that a complete description should afford. One of the largest discrepancies occurs at the Schiff base nitrogen site, where the predicted shift is greater than three standard errors from the experimental value.

**Fig. 5.**
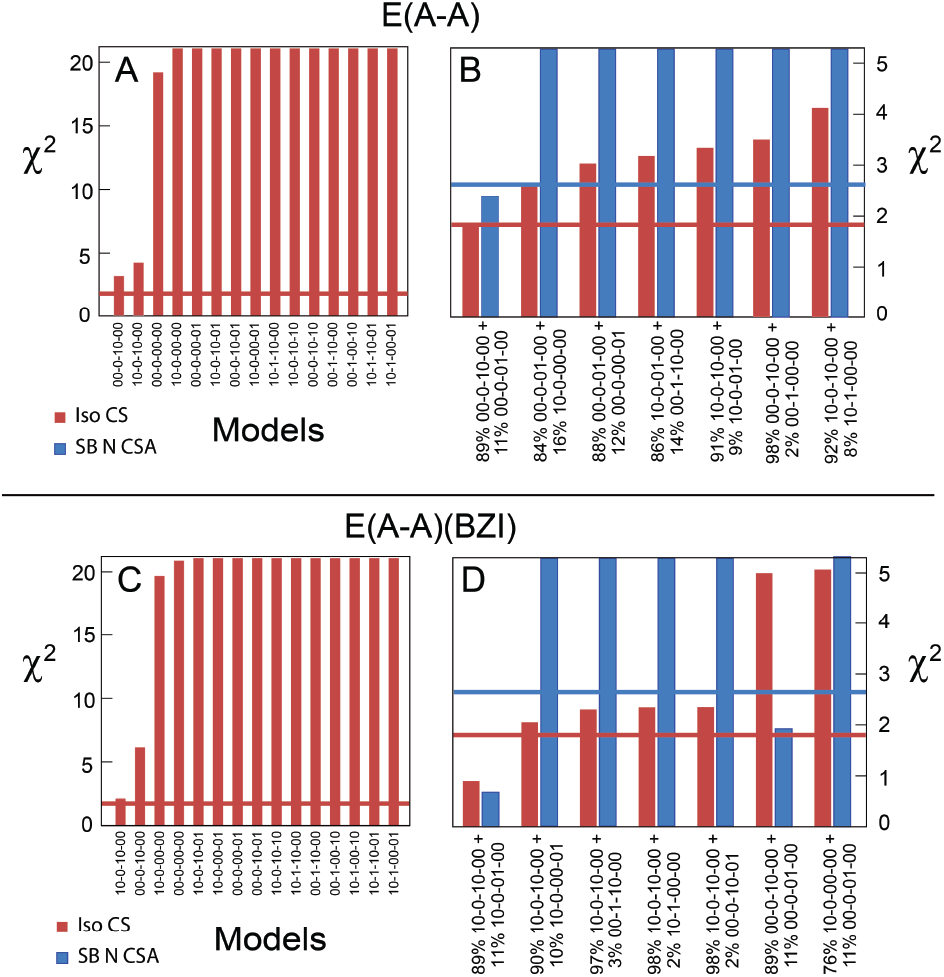
Ranking of structural models based on agreement between the experimental and first principles chemical shifts. (A,C) The 15 best geometry-optimized active site models for (A) the E(A-A) intermediate and (C) the E(A-A)(BZI) complex; structures and labeling given in Schemes S1 and S2. (B, D) Rankings of the 7 best fastexchange equilibrium models comparing the experimental and first-principles isotropic (red) and Schiff base nitrogen chemical shift tensor components (blue). 95% confidence limits are shown as the correspondingly colored horizontal bars.

Motivated by the temperature dependence of the Schiff base nitrogen chemical shift, a simple two-site, fast-exchange equilibrium model was considered next in which the effective chemical shift was given as the population-weighted average of the individual shifts. To remain unbiased, all structures that differed by the position of a single hydrogen atom were paired and their populations optimized for best overall agreement with the experimental chemical shifts. These models were again ranked and are summarized in Fig. 5. Multiple models satisfy or come close to satisfying the 95% confidence limits for the chemical shift reducued-χ^2^. To delineate these models, their predicted Schiff base nitrogen chemical shift tensors were compared to the experimental tensor. The chemical shift tensor is extremely sensitive to chemical exchange as its components average not only in magnitude, but also in orientation. As shown in Fig. 5, only a single tautomeric equilibrium model for both E(A-A) and E(A-A)(BZI) simultaneously satisfy the isotropic and anisotropic chemical shift restraints, indicating that the benchmark for a solved structure has been reached in both cases. For both E(A-A) and E(A-A)(BZI), the best-fit equilibrium is between the phenolic (89%) and protonated Schiff base (11%) tautomers, with proton exchange across the internal hydrogen bond (Table 1, center inset); this exchange is analogous to that found for the carbanionic intermediate (23) and appears to be a common feature in PLP-dependent enzymes (8). The only difference in protonation states between E(A-A) and E(A-A)(BZI) is for the ε-amino group of βLys87: for E(A-A) it is neutral, while for E(A-A)(BZI) it is positively charged.

As a final test of consistency for the exchange model, the ^15^N chemical shifts of the Schiff base nitrogen in both structures were measured at 95 K under conditions of dynamic nuclear polarization as shown in Fig. S5 and detailed in the SI. At 95 K, the tautomeric equilibrium is expected to shift predominantly to the phenolic form, with less than 1% of the exchange partner present. This is confirmed experimentally, with chemical shifts of 301.4 ppm for E(A-A) and 302.1 ppm for E(A-A)(BZI) at 95 K, compared to the predicted shifts for the individual major structures of 302.9 ppm and 310.2 ppm, respectively. Thus, both the major tautomers and the dynamic equilibrium of these intermediates can be established through NMR-assisted crystallography.

### Chemically-Rich Crystal Structures and Mechanistic Implications

What emerges from the application of NMR-assisted crystallography to the E(A-A) intermediate and the E(A-A)(BZI) complex in TS is a remarkable, chemically-detailed view of the enzyme active sites as shown in Fig. 6. In these models, not only are the heavy atom locations from X-ray crystallography shown, but the hydrogen atom locations have been determined. While the pictures in Fig. 6 are static, the systems are not: the microcrystalline samples remain catalytically active and dynamic. We interpret these structures as equilibrium geometries of the intermediates, about which fluctuations do occur. For tautomeric exchange, the fluctuations are large enough that they must be accounted for in the first-principles analysis. For the other motions, the deviations are presumably small enough that the average structure serves as a reasonable proxy. Considerable dynamics of the waters proximal to the substrate may explain why the overall agreement between theory and experiment is lower for E(A-A).

**Fig. 6:**
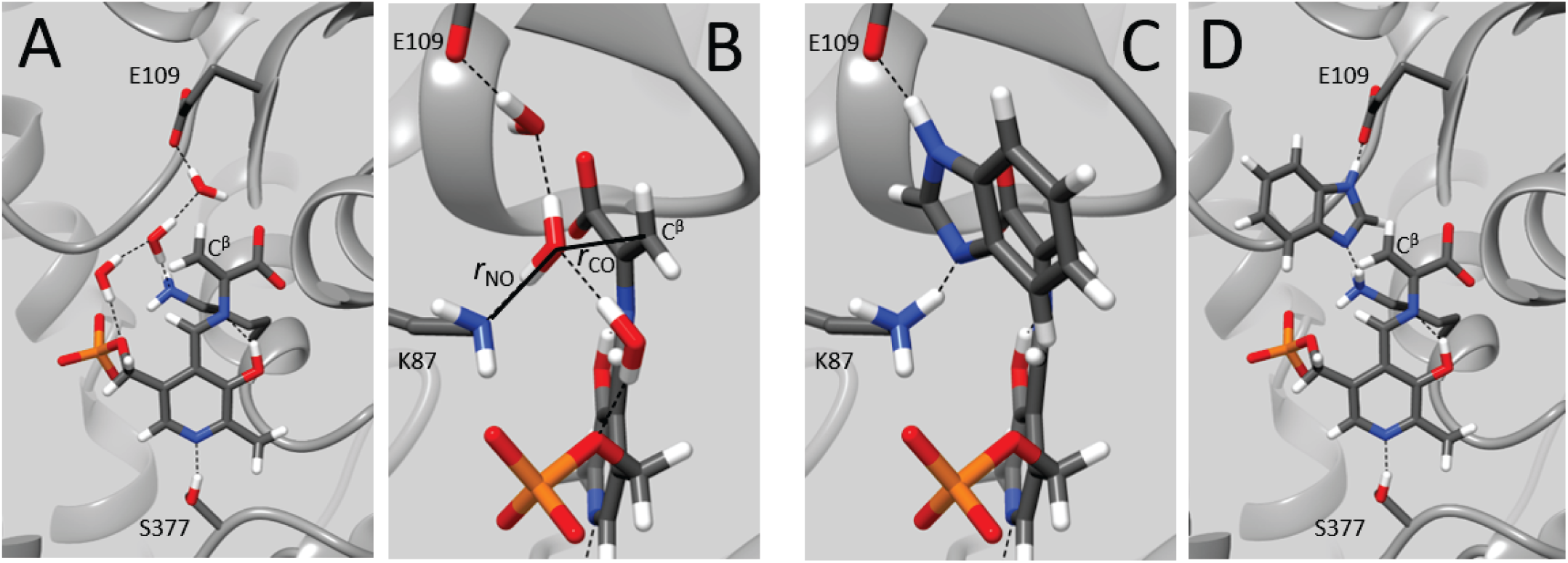
Protonation states, hydrogen bonding interactions, and the placement and orientation of structural waters as revealed by NMR-assisted crystallography in the tryptophan synthase β-subunit active site. (A,B) the E(A-A) intermediate and (C,D) the E(A-A)(BZI) complex. For E(A-A), the position and orientation of the center water (B) suggests the pathway back to the transition state for the loss of the serine β-hydroxyl and the identification of βLys87 as the active site acid-base catalytic residue. The E(A-A)(BZI) complex shows BZI bound in the active site with hydrogen bonding interactions to βGlu109 and the charged ε-amino group of βLys87. BZI is poised for, but unable to initiate, the subsequent bond formation step. Images rendered in UCSF Chimera (4).

Perhaps most striking for E(A-A) is the orientation of the central water molecule immediately adjacent to the serine substrate C^β^ and positioned with one hydrogen pointing back toward the ε-amino group of the active site acid-base catalytic residue βLys87 (Fig. 6B). This water is located 2.8 Å distant from the C^β^ and is perfectly aligned for the reverse nucleophilic attack at C^β^ to regenerate E(C1). The interaction of the βLys87 ε-NH2 group with this water and its proximity to the C^β^ carbon support the hypothesis that βLys87 is the acid catalyst that protonates the β-hydroxyl in the transition state for C-O bond scission as E(C1) is converted to E(A-A). It is remarkable that the placement and orientation of this water suggests that it is only tenths of Angstroms away from the preceding transition state with an orientation that points along the reaction coordinate.

Critical examination of the crystal structures for the E(A-A) intermediate, however, shows that this central water is not consistently present. Scrutiny of the electron density maps for 2J9X and 4HN4 (both with Cs^+^ in the monovalent cation binding site) show significantly lower electron density for the center water, and likely reflects occupancy lower than 50% (Fig. S10). To explore this variation, we prepared and solved two additional E(A-A) crystal structures at 1.40 Å and 1.50 Å, respectively, for the NH✓ and Cs^+^ forms (PDBID’s 7MT4 and 7MT5). The diffraction data (summarized in Table S4) show well-defined electron density for the central water in the NH4^+^ form, but a complete lack of this density in the Cs^+^ structure (Fig. S10). Close examination of the NMR data for E(A-A) compared to E(A-A)(BZI) indicates that each of the ^13^C resonances that derive from the substrate serine is accompanied by slightly shifted (±1-2 ppm) minor species of approximately 20-30% intensity, while those for E(A-A)(BZI) show no such peaks. These resonances are most easily observed when single site ^13^C labels are incorporated on serine, as shown in Fig. 7A, and display the same multiplet pattern as the major resonances upon incorporation of 2,3-^13^C2-L-Ser. The minor and major resonances also show distinct sets of cross-peaks (minor to minor, major to major) in the 2D dipolar-driven CORD correlation experiment (53) (Fig. 7C), acquired here with a CPMAS cryoprobe to improve sensitivity (3). Motivated by the variability in the placement of the central water in the crystal structures, we hypothesize that the sets of major and minor resonances belong to two independent E(A-A) species that derive from varying occupancy of the center water across the macroscopic crystals. To test this, we constructed active site models in which the water nearest C^β^ was removed for the primary tautomers identified above. The structures were fully geometry optimized and their chemical shifts again predicted using DFT (Table S1). The predicted chemical shifts nicely track the experimental chemical shifts of the minor species, and we tentatively assign these peaks to a minor population in which the central water is no longer bound. Taken together, the solid-state NMR, X-ray, and computational results are consistent with this site being a weak binding site for the water and highlight the incredible sensitivity to the active site chemistry that can be obtained through this synergistic and self-consistent approach.

**Fig. 7:**
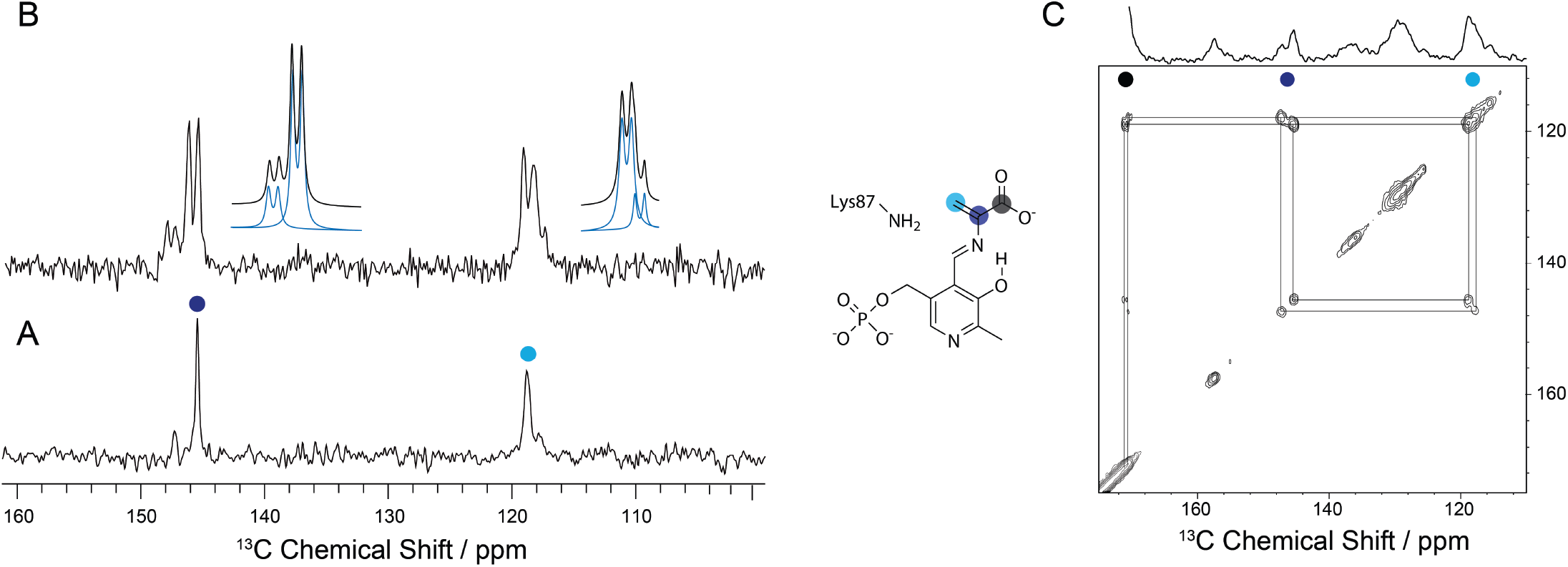
^13^C SSNMR CPMAS spectra of the microcrystalline TS E(A-A) intermediate prepared with (A) a 50:50 mixture of singly-labeled 2-^13^C-L-Ser and singly-labeled 3-^13^C-L-Ser and (B) doubly labeled 2,3-^13^¤2-L-Ser. For both, the natural abundance ^13^C protein background has been subtracted as in Fig. 3. The C^α^ and C^β^ resonances show a minor peak that, along with the major peak, splits into a doublet upon incorporation of doubly labeled L-Ser substrate (deconvolution shown adjacent to the spectrum). (C) The 2D dipolar driven ^13^C correlation spectrum of the E(A-A) formed with U-^13^C3-L-Ser displays distinct crosspeaks for the major and minor peaks, indicating that they belong to two independent E(A-A) species. The small chemical shift changes for the minor peaks correlate with the first-principles predicted shifts in the absence of the center water in E(A-A), consistent with the variable occupancy of this site across X-ray crystal structures. 1D spectra acquired as in Fig. 3; 2D Spectra acquired using CORD mixing on a Bruker BioSolids CryoProbe^™^ (3) at 14.1 T, −10 °C (sample temp), and 8 kHz MAS as described in the Methods and Materials (SI).

The E(A-A)(BZI) complex shows a similarly remarkable view of the active site chemistry, but now for the subsequent mechanistic step of bond formation between C^β^ and the incoming nucleophile. BZI is an isostere of indole and a potent inhibitor of the β-reaction (35, 38, 54, 55). The view from NMR-assisted crystallography (Fig. 6C,D) shows BZI bound in the active site, displacing the three neighboring crystallographic waters, and poised for nucleophilic attack on C^β^. Despite being a much better nucleophile than indole, BZI does not react to form a covalent bond. A comparison to crystal structures for the carbanionic intermediate analogs formed with indoline and 2-aminophenol (23) makes it clear that BZI is bound to the same subsite. Thus, this intermediate analog appears to be poised within a few tenths of an Angstrom of the transition state between two reactive intermediates in the TS catalytic cycle.

That BZI is not a substrate for TS and does not covalently react with E(A-A) can be attributed to two important factors: the different nucleophilic reaction mechanisms of indole and BZI, and the tight packing and hydrogen bonding interactions within the indole subsite. The mechanisms of the nucleophilic reactions of indole and BZI are fundamentally distinct (54). Each nitrogen of BZI formally has an unshared electron pair, but these nitrogens are very different. The unshared electron pair on N1 resides in a 2p orbital that is integral to the π electron aromatic system. This nitrogen also has a proton covalently bonded to it. This nitrogen is not a target for bonding to electrophiles. The electron pair of the other nitrogen (N3) is not part of the aromatic π system and resides in an sp^2^ hybrid orbital oriented orthogonal to the BZI π-system. It is this pair of electrons that functions as a nucleophile in BZI reactions. As shown in Fig. 6C,D, NMR-assisted crystallography identifies that it is the nucleophilic N3 that is adjacent to the substrate C^β^ in the E(A-A)(BZI) complex. Importantly, the structure shows that the BZI sp^2^ hybrid orbital containing the nucleophilic electron pair does not point toward the E(A-A) C^β^ p-orbital. The tight packing of atoms from the protein and PLP at the indole subsite, the hydrogen bond between N3 and the protonated ε-amino group nitrogen of βLys87, and the hydrogen bond between the βGlu109 carboxylate and N1 of BZI preclude rearrangement of BZI within the site to allow the reaction.

When indole is modeled in place of BZI in the E(A-A)(BZI) complex, the C3 carbon of the 5-membered ring is perfectly aligned to form the new C-C bond. The attacking p-orbital points toward the electron deficient C^β^ carbon and is poised to make orbital overlap as the complex moves along the reaction coordinate to the sp^3^ geometry of the transition state and the product. At the same time, N1 of indole is positioned to hydrogen bond to the carboxylate of βGlu109 and stabilize positive charge development on N1 as the transition state is approached. This turns out to be critical from a mechanistic standpoint. Owing to its enamine structure, the C3 carbon of indole is relatively electron rich; nevertheless, indole is a poor nucleophile and requires assistance in the attack on C^β^ of the α-aminoacrylate intermediate. This assistance is provided by βGlu109. Indeed, the βGlu109Asp mutation reduces the β reaction rate 27-fold, while reaction of the indole analogue indoline to give the L-Trp analogue dihydroiso-L-tryptophan is increased compared to WT TS (39). Indoline is a relatively good nucleophile and does not require the assistance of βGlu109. The chemically-detailed, three-dimensional structure of the E(A-A)(BZI) complex (Fig. 6D) establishes that the BZI N1 is positioned to hydrogen bond to the carboxylate of βGlu109. In the corresponding indole complex this interaction would stabilize positive charge development on N1 during the nucleophilic attack. We posit that the E(A-A)(BZI) complex models how indole is bound to E(A-A) just prior to formation of the transition state for C-C bond formation.

## Conclusions

The joint application of solid-state NMR spectroscopy, X-ray crystallography, and first principles computational chemistry offers unprecedented, chemically-rich views of the TS E(A-A) and E(A-A)(BZI) active sites. Through the combined determination of heavy atom positions, the unique identification of protonation states, and the placement and orientation of active site waters, an exceptionally clear picture of structure and reactivity emerges. Remarkably, both E(A-A) and E(A-A)(BZI) appear poised near, but unable to initiate, chemical reaction. The E(A-A) intermediate shows a central water that is well positioned for the reverse nucleophilic attack on C^β^, with an orientation that points back to the active site acid-base catalytic residue, βLys87, highlighting the reaction coordinate for the elimination of the substrate β-hydroxyl. Both X-ray crystallography and solid-state NMR suggest variable occupancy for this site, underscoring the chemical sensitivity of the approach. Upon reaction of E(A-A) with BZI, these waters are displaced, and NMR-assisted crystallography shows BZI occupying the presumed binding pocket for the natural substrate, indole, but again unable to react. Here the protonation states complete the chemical picture for why BZI, despite being a good nucleophile, is unable to initiate the next step in the chemical reaction, as it is held in the wrong orientation by hydrogen bonds to βGlu109 and the charged ε-amino group of βLys87.

NMR crystallography takes advantage of one of the well-established strengths of NMR spectroscopy – remarkable sensitivity to chemical structure and chemical dynamics. This is, we would argue, where NMR will interface most strongly with the other tools of structural biology, including X-ray and neutron crystallography and Cryo-EM. When combined with first-principles computational chemistry, these complementary techniques can build consistent, testable models of structure and reactivity in enzyme active sites. Remarkably, this is accomplished here near room temperature and under conditions of active catalysis.

## Data Availability

All new crystal structures reported here have been deposited with the Protein Data Bank. All NMR chemical shift data are reported in the article and/or supporting information. Any additional data or processing scripts are freely available from the authors upon request.

## Acknowledgements

The PI’s thank Professor Martin Safo for providing the ePL Kinase K229Q mutant (56).

Research reported in this paper was supported by the United States National Science Foundation (CHE-1710671 to LJM, NSF-1708019 to RJH) and the US National Institutes of Health (GM097569 to LJM, MFD; GM137008 to LJM and TCM; NIH P41 GM122698 to JRL). Computations were performed using the computer clusters and data storage resources of the UCR HPCC, which were funded by grants from NSF (MRI-1429826) and NIH (S10OD016290). A portion of this work was performed at the National High Magnetic Field Laboratory, which is supported by National Science Foundation Cooperative Agreement Nos. DMR-1157490 and DMR-1644779 and the State of Florida. The gyrotron was purchased through an NSF MRI award, CHE-1229170, and the magnet and NMR console were purchased through NIH S10 OD018519. Development of the SCH magnet and NMR instrumentation were supported by NSF instrumentation grants (DMR-1039938 and DMR-0603042).

## Materials and Methods

Protein crystals of E(A-A) and E(A-A)(BZI) were prepared and structures solved following our earlier protocols (35, 57). First principles calculations were performed using a cluster-based model of the active site as described previously (23). Tryptophan synthase was prepared by overexpression of *St*TS in *E. coli* (40, 44) and used for SSNMR and X-ray crystallography as described earlier (23, 57). SSNMR experiments followed our prior experimental design (23, 46), including DNP experiments at 95 K (58), and 2D correlation experiments performed on a cryoMAS probe (3). ^15^N-benzimidazole (BZI) was synthesized from ^15^NH_4_OH and 1-fluoro-2-nitrobenzene as described in the SI; 2,2’,3-^13^C_3_,^15^N-PLP was prepared from U-^13^C_3_,^15^N-Ala as previously detailed (44), while ^15^N-PLP was prepared with a new synthetic strategy including enzymatic phosphorylation via the ePL Kinase K229Q mutant (56). Detailed Materials and Methods are included as part of the Supporting Information.

## Notes

**Competing Interest Statement:**

### Competing Interest Statement

The authors have declared no competing interest.

## References

1. M. F. Dunn, Allosteric regulation of substrate channeling and catalysis in the tryptophan synthase bienzyme complex. Arch. Biochem. Biophys. 519, 154–166 (2012).

2. E. W. Miles, Stereochemistry and mechanism of a new single-turnover, half-transamination reaction catalyzed by the tryptophan synthase alpha 2 beta 2 complex. Biochemistry 26, 597–603 (1987).

3. A. Hassan et al., Sensitivity boosts by the CPMAS CryoProbe for challenging biological assemblies. J. Magn. Reson. 311, 106680 (2020).

4. E. F. Pettersen et al., UCSF Chimera--a visualization system for exploratory research and analysis. J Comput Chem 25, 1605–1612 (2004).

5. C. Walsh, Enzymatic Reaction Mechanisms (W. H. Freeman and Company, San Francisco, 1979), pp. 978.

6. H. Hayashi, Pyridoxal enzymes: mechanistic diversity and uniformity. J. Biochem. 118, 463–473 (1995).

7. R. Percudani, A. Peracchi, A genomic overview of pyridoxal-phosphate-dependent enzymes. EMBO Rep 4, 850–854 (2003).

8. M. D. Toney, Controlling reaction specificity in pyridoxal phosphate enzymes. Biochim Biophys Acta 1814, 1407–1418 (2011).

9. E. W. Miles, Tryptophan synthase: structure, function, and subunit interaction. Adv. Enzymol. Relat. Areas Mol. Biol. 49, 127–186 (1979).

10. C. Yanofsky, I. P. Crawford, “Tryptophan Synthetase” in The Enzymes Third Edition, P. D. Boyer, Ed. (Academic Press, Inc., New York, NY, 1972), vol. 7, chap. 1, pp. 1–32.

11. M. F. Dunn, D. Niks, H. Ngo, T. R. Barends, I. Schlichting, Tryptophan synthase: the workings of a channeling nanomachine. Trends Biochem Sci 33, 254–264 (2008).

12. M. Chan-Huot, S. Sharif, P. M. Tolstoy, M. D. Toney, H. H. Limbach, NMR Studies of the Stability, Protonation States, and Tautomerism of 13C- and ^15^N-Labeled Aldimines of the Coenzyme Pyridoxal 5’-Phosphate in Water. Biochemistry 49, 10818–10830 (2010).

13. M. Chan-Huot et al., NMR studies of protonation and hydrogen bond states of internal aldimines of pyridoxal 5’-phosphate acid-base in alanine racemase, aspartate aminotransferase, and poly-L-lysine. J Am Chem Soc 135, 18160–18175 (2013).

14. P. S. Langan et al., Substrate binding induces conformational changes in a class A β-lactamase that prime it for catalysis. ACS Catalysis 8, 2428–2437 (2018).

15. S. Dajnowicz et al., Direct visualization of critical hydrogen atoms in a pyridoxal 5’-phosphate enzyme. Nat Commun 8, 955 (2017).

16. K. M. Yip, N. Fischer, E. Paknia, A. Chari, H. Stark, Atomic-resolution protein structure determination by cryo-EM. Nature 587, 157–161 (2020).

17. D. F. Gauto et al., Integrated NMR and cryo-EM atomic-resolution structure determination of a half-megadalton enzyme complex. Nat Commun 10, 2697 (2019).

18. R. Michalczyk et al., Joint neutron crystallographic and NMR solution studies of Tyr residue ionization and hydrogen bonding: Implications for enzyme-mediated proton transfer. Proc Natl Acad Sci U S A 112, 5673–5678 (2015).

19. A. Carlon et al., Improved Accuracy from Joint X-ray and NMR Refinement of a Protein-RNA Complex Structure. J Am Chem Soc 138, 1601–1610 (2016).

20. K. Zhang et al., Structure of the 30 kDa HIV-1 RNA Dimerization Signal by a Hybrid Cryo-EM, NMR, and Molecular Dynamics Approach. Structure 26, 490–498 e493 (2018).

21. J. Lai et al., X-ray and NMR crystallography in an enzyme active site: the indoline quinonoid intermediate in tryptophan synthase. J Am Chem Soc 133, 4–7 (2011).

22. L. J. Mueller, M. F. Dunn, NMR crystallography of enzyme active sites: probing chemically detailed, threedimensional structure in tryptophan synthase. Acc. Chem. Res. 46, 2008–2017 (2013).

23. B. G. Caulkins et al., NMR Crystallography of a Carbanionic Intermediate in Tryptophan Synthase: Chemical Structure, Tautomerization, and Reaction Specificity. J Am Chem Soc 138, 15214–15226 (2016).

24. J. C. Facelli, D. M. Grant, Determination of molecular symmetry in crystalline naphthalene using solid-state NMR. Nature 365, 325–327 (1993).

25. C. Bonhomme et al., First-principles calculation of NMR parameters using the gauge including projector augmented wave method: a chemist’s point of view. Chem Rev 112, 5733–5779 (2012).

26. M. Baias et al., De novo determination of the crystal structure of a large drug molecule by crystal structure prediction-based powder NMR crystallography. Journal of the American Chemical Society 135, 17501–17507 (2013).

27. D. H. Brouwer et al., A general protocol for determining the structures of molecularly ordered but noncrystalline silicate frameworks. J Am Chem Soc 135, 5641–5655 (2013).

28. C. Martineau, NMR crystallography: Applications to inorganic materials. Solid State Nucl. Magn. Reson. 63-64, 1-12 (2014).

29. C. Luchinat, G. Parigi, E. Ravera, M. Rinaldelli, Solid-state NMR crystallography through paramagnetic restraints. J Am Chem Soc 134, 5006–5009 (2012).

30. A. Bertarello et al., Picometer Resolution Structure of the Coordination Sphere in the Metal-Binding Site in a Metalloprotein by NMR. J Am Chem Soc 142, 16757–16765 (2020).

31. H. Zhang et al., HIV-1 Capsid Function Is Regulated by Dynamics: Quantitative Atomic-Resolution Insights by Integrating Magic-Angle-Spinning NMR, QM/MM, and MD. J Am Chem Soc 138, 14066–14075 (2016).

32. C. H. Tai, P. F. Cook, O-acetylserine sulfhydrylase. Adv. Enzymol. Relat. Areas Mol. Biol. 74, 185–234 (2000).

33. R. S. Phillips et al., Pressure and Temperature Effects on the Formation of Aminoacrylate Intermediates of Tyrosine Phenol-lyase Demonstrate Reaction Dynamics. ACS Catalysis 10, 1692–1703 (2019).

34. H. Ngo et al., Allosteric regulation of substrate channeling in tryptophan synthase: modulation of the L-serine reaction in stage I of the beta-reaction by alpha-site ligands. Biochemistry 46, 7740–7753 (2007).

35. D. Niks et al., Allostery and substrate channeling in the tryptophan synthase bienzyme complex: evidence for two subunit conformations and four quaternary states. Biochemistry 52, 6396–6411 (2013).

36. M. A. Maria-Solano, J. Iglesias-Fernandez, S. Osuna, Deciphering the Allosterically Driven Conformational Ensemble in Tryptophan Synthase Evolution. J Am Chem Soc 141, 13049–13056 (2019).

37. A. R. Buller et al., Directed Evolution Mimics Allosteric Activation by Stepwise Tuning of the Conformational Ensemble. J Am Chem Soc 140, 7256–7266 (2018).

38. M. F. Dunn et al., The tryptophan synthase bienzyme complex transfers indole between the alpha- and beta-sites via a 25-30 A long tunnel. Biochemistry 29, 8598–8607 (1990).

39. P. S. Brzovic, A. M. Kayastha, E. W. Miles, M. F. Dunn, Substitution of glutamic acid 109 by aspartic acid alters the substrate specificity and catalytic activity of the beta-subunit in the tryptophan synthase bienzyme complex from Salmonella typhimurium. Biochemistry 31, 1180–1190 (1992).

40. B. G. Caulkins et al., Catalytic roles of betaLys87 in tryptophan synthase: (15)N solid state NMR studies. Biochim Biophys Acta 1854, 1194–1199 (2015).

41. L. M. Mcdowell, M. S. Lee, J. Schaefer, K. S. Anderson, Observation of an Aminoacrylate Enzyme Intermediate in the Tryptophan Synthase Reaction by Solid-State Nmr. Journal of the American Chemical Society 117, 12352–12353 (1995).

42. J. Herzfeld, A. E. Berger, Sideband Intensities in Nmr-Spectra of Samples Spinning at the Magic Angle. Journal of Chemical Physics 73, 6021–6030 (1980).

43. V. Copie, W. S. Faraci, C. T. Walsh, R. G. Griffin, Inhibition of alanine racemase by alanine phosphonate: detection of an imine linkage to pyridoxal 5’-phosphate in the enzyme-inhibitor complex by solid-state nitrogen-15 nuclear magnetic resonance. Biochemistry 27, 4966–4970 (1988).

44. B. G. Caulkins et al., Protonation states of the tryptophan synthase internal aldimine active site from solid-state NMR spectroscopy: direct observation of the protonated Schiff base linkage to pyridoxal-5’-phosphate. J Am Chem Soc 136, 12824–12827 (2014).

45. K. D. Schnackerz, B. Andi, P. F. Cook, (31)P NMR spectroscopy senses the microenvironment of the 5’-phosphate group of enzyme-bound pyridoxal 5’-phosphate. Biochim Biophys Acta 1814, 1447–1458 (2011).

46. R. P. Young et al., Solution-State 17O Quadrupole Central-Transition NMR Spectroscopy in the Active Site of Tryptophan Synthase. Angewandte Chemie International Edition 55, 1350–1354 (2016).

47. J. Zhu, G. Wu, Quadrupole central transition 17O NMR spectroscopy of biological macromolecules in aqueous solution. J Am Chem Soc 133, 920–932 (2011).

48. Z. Gan et al., NMR spectroscopy up to 35.2T using a series-connected hybrid magnet. J. Magn. Reson. 284, 125–136 (2017).

49. T. Gullion, J. Schaefer, Rotational-Echo Double-Resonance Nmr. Journal of Magnetic Resonance 81, 196–200 (1989).

50. J. D. Hartman, T. J. Neubauer, B. G. Caulkins, L. J. Mueller, G. J. Beran, Converging nuclear magnetic shielding calculations with respect to basis and system size in protein systems. J. Biomol. NMR 62, 327–340 (2015).

51. J. D. Hartman, R. A. Kudla, G. M. Day, L. J. Mueller, G. J. Beran, Benchmark fragment-based (1)H, (13)C, (15)N and (17)O chemical shift predictions in molecular crystals. Phys. Chem. Chem. Phys. 18, 21686–21709 (2016).

52. S. T. Holmes, R. J. Iuliucci, K. T. Mueller, C. Dybowski, Critical Analysis of Cluster Models and Exchange-Correlation Functionals for Calculating Magnetic Shielding in Molecular Solids. J Chem Theory Comput 11, 5229–5241 (2015).

53. G. Hou, S. Yan, J. Trebosc, J. P. Amoureux, T. Polenova, Broadband homonuclear correlation spectroscopy driven by combined R2(n)(v) sequences under fast magic angle spinning for NMR structural analysis of organic and biological solids. J. Magn. Reson. 232, 18–30 (2013).

54. M. Roy, S. Keblawi, M. F. Dunn, Stereoelectronic control of bond formation in Escherichia coli tryptophan synthase: substrate specificity and enzymatic synthesis of the novel amino acid dihydroisotryptophan. Biochemistry 27, 6698–6704 (1988).

55. R. M. Harris, H. Ngo, M. F. Dunn, Synergistic effects on escape of a ligand from the closed tryptophan synthase bienzyme complex. Biochemistry 44, 16886–16895 (2005).

56. M. S. Ghatge et al., Pyridoxal 5’-phosphate is a slow tight binding inhibitor of E. coli pyridoxal kinase. PLoS One 7, e41680 (2012).

57. E. Hilario et al., Visualizing the tunnel in tryptophan synthase with crystallography: Insights into a selective filter for accommodating indole and rejecting water. Biochim Biophys Acta 1864, 268–279 (2016).

58. X. Wang et al., Direct dynamic nuclear polarization of 15N and 13C spins at 14.1 T using a trityl radical and magic angle spinning. Solid state nuclear magnetic resonance 100, 85–91 (2019).

