## Supporting Information for "Imaging active site chemistry and protonation states: NMR crystallography of the tryptophan synthase α-aminoacrylate intermediate"

**[1]co-first authors**

**This PDF file includes:**

Supplementary text: Expanded Materials and Methods
Figures S1 to S10
Tables S1 to S4
Schemes S1 to S2
SI References

### Materials and Methods

First-Principles Calculations

First principles calculations were performed using a cluster-based model of the active site as described previously (1). The clusters for the TS E(A-A) and E(A-A)(BZI) were constructed from the corresponding crystal structures 4HN4 and 4HPX by selecting all atoms within 7 Å of the cofactor-substrate complex (Fig. S1A,B). This selection was expanded to include complete residues and were modified as follows: (1) residues that were not part of continuous backbone segments and with only two atoms within the initial 7 Å cut were deleted (Q142, V201, I238, D383); (2) residues with two backbone atoms and no side chain residues within 7 Å were converted to alanine (N145, K167, D305, S308, V309, L349, N375, L376, R379); (3) residues with two side chain atoms and no backbone atoms within 7 Å were truncated by removing the backbone atoms and replacing $C^{\alpha}$ with a methyl group (β-site residues F280, H313, K382); (4) N-terminal nitrogen atoms were replaced with a hydrogen atom and C-terminal carbonyls were capped with an $-NH_2$ group (amidated); (5) the $Cs^{+}$ cation was replaced with $Na^{+}$; and (6) hydrogen atoms were added. The final model for E(A-A) included 58 amino acid residues, the PLP-Serine complex, and 13 water molecules (waters 511, 515, 516, 519, 535, 538, 559, 570, 583, 604, 744, 844, 914) and had a total of 703 atoms. The final model for E(A-A)(BZI) included the same residues, but also included the BZI inhibitor and only 10 water molecules (HOH 505, 511, 513, 516, 520, 541, 549, 683, 688, and 710). The structures were further modified depending on the protonation states of the cofactor-substrate complex and K87.

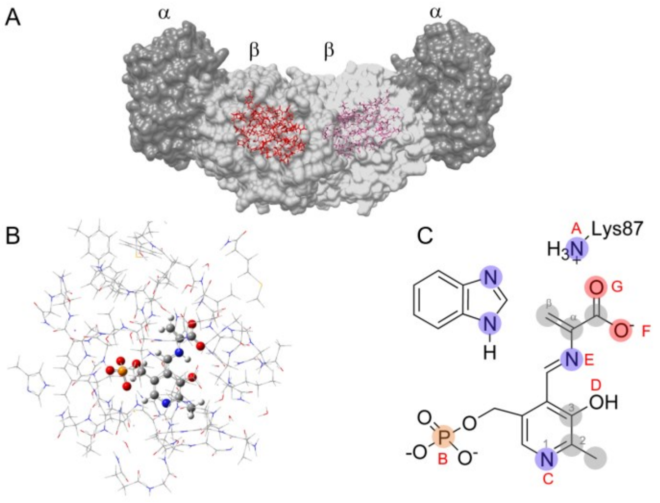


Fig. S1 (repeated from Fig. 4, main text). (A) X-ray crystal structure (PDBID: 4HN4) of the tryptophan synthase $\alpha_2\beta_2$ heterodimer with the β-subunit active site in red. (B) Cluster model of the active site for first-principles geometry optimization and chemical shift calculations with the protein side chains shown in wireframe and the cofactor and substrates in ball-and-stick. (C) Protonation sites on and near the cofactor/substrate complex: A the βLys87 side chain, B the PLP phosphate group, C the PLP pyridine ring nitrogen, D the PLP phenolic oxygen, E the Schiff-base nitrogen, and F,G the substrate carboxylate. Shaded nuclei indicate sites for which experimental NMR chemical shifts

Within this cluster, candidate structures were systematically generated by varying the protonation of the following seven ionizable sites on or near the PLP-ligand complex (Fig. S1C): the ε-amino group of βLys87, the PLP phosphate group, the pyridine nitrogen, the pyridoxal oxygen, the Schiff base nitrogen, and both carboxylate oxygen atoms; the protonation states of the two BZI nitrogen atoms were also changed for the relevant structures. For both E(A-A) and E(A-A)(BZI), the phosphate group was taken to be dianionic based on its CSA tensor, and models that had more than a single proton placed at either the pyridoxal oxygen or the Schiff base nitrogen were not considered, nor were structures with a doubly protonated carboxylate. Based on its $^{15}N$ Chemical shifts, BZI was modeled as neutral as shown in Fig. S1C and oriented with the protonated nitrogen adjacent to βGlu109. Initial structures with the protonated nitrogen directed toward a neutral βLys87 with βGlu109 protonated invariably resulted in a redistribution of protonation states during geometry optimization to give protonated βLys87 and ionized βGlu109. In sum, 35 variations of protonation state were constructed for E(A-A) and E(A-A)(BZI). These models were labeled using a binary code to indicate whether a site was protonated ("1" – yes, "0" – no) in the following order: (Nε of βLys87)(phosphate group)-(pyridine nitrogen)-(pyridoxyl phenolic oxygen)(Schiff base nitrogen)-(nearer carboxylate oxygen to the Schiff base)(farther carboxylate oxygen); these sites are designated as AB-C-DE-FG in Fig. S1 and shown explicitly in Schemes S1 and S2.

Initial proton orientations in the active site clusters were set using molecular dynamics (MD) simulations of the αβ dimer. These simulations were performed using Amber package with GPU acceleration (2, 3). Coordinates for *StTS* were obtained from crystal structure with PDB ID 4HN4 for E(A-A) and 4HPX for AA/BZI with crystal waters retained (4). Amber force field FF14SB was applied to protein atoms (5). Ligands were parametrized using General Amber force field (GAFF) and AM1-bcc charge model (6, 7). Only hydrogens were minimized and the MD simulations were carried out using Generalized Born implicit solvent model (GB-Neck2)(8) at constant temperature of 298 K controlled by Langevin thermostat. Two MD trajectories for each system were collected over 50 ns at 1 ps interval with 2 fs timestep. The trajectories were further processed with CPPTRAJ software to contain 5000 frames for analysis (9). The systems were visualized and analyzed using Visual Molecular Dynamics software (10). Dihedral data was collected for residues Thr110, Thr190, Ser235, Ser377, Ser351 and Tyr186 in both systems. In E(A-A) bound system (4HN4), the conformation and orientation for three water molecules (WAT583, WAT604, WAT914) in beta subunit were analyzed. The initial conformation and the position of the polar hydrogens in the respective residues and water molecules were selected based on the mean value of the normally distributed dihedral data for each residue.

All cluster models were subsequently geometry optimized in Gaussian09 (11) at the DFT B3LYP level of theory using the Grimme D3 empirical dispersion correction (12) and a two-tier, locally-dense basis set (13-15) with 6-31G(d,p) for the PLP/substrate complex and 6-31G for all other atoms. Except as noted below, all atoms on or within 4 Å of the PLP/substrate complex and all hydrogen atoms in the cluster could adjust while the remaining atom coordinates were fixed at their crystallographic values. The only exceptions were the ε-amino group nitrogen of βLys87 and the oxygen atoms of the three waters adjacent to the serine moiety for the α-aminoacrylate, which were also held fixed at their crystallographic coordinates during the geometry optimizations.

For each optimized structure, NMR shieldings were calculated at the DFT B3LYP level of theory and employing a three-tier, locally dense basis set assignment with the PLP/substrate, βLys87, BZI, and proximal water atoms at 6-311+G(2d,p), atoms within 4Å of this subgroup at 6-311G(d,p), and all remaining atoms at 6-31G. NMR shielding values were converted to chemical shifts using the following linear rescaling relationships, derived at the same level of theory and bases (13, 14).

$$^{13}\text{C:}\quad \delta\left[\text{TMS}(l)\right] = 173.70 - 0.9685\,\sigma_{\text{calc}} \tag{1}$$

$$^{15}\text{N:}\quad \delta\left[\text{NH}_3(l)\right] = \delta\left[\text{NH}_4\text{Cl}(s)\right] + 39.27 = 230.45 - 0.9996\,\sigma_{\text{calc}} \tag{2}$$

$$^{17}\text{O:}\quad \delta\left[\text{H}_2\text{O}(l)\right] = 266.32 - 1.0551\,\sigma_{\text{calc}} \tag{3}$$

Previous benchmark studies across test sets of solid-state structures used to establish these relations, demonstrated root-mean-square errors (RMSE) for isotropic shifts to be 1.5 ppm for $^{13}$C, 4.3 ppm for $^{15}$N, and 7.5 ppm for $^{17}$O (13). RMSE for the chemical shift tensor components were 4.2 ppm and 13.7 ppm for $^{13}$C and $^{15}$N respectively (1).

The structural models were ranked using the reduced-$\chi^2$ statistic,(16) a quantitative measure of the agreement between their first-principles predicted chemical shifts and the experimental NMR parameters,

$$\chi_r^2(\text{model}) = \frac{1}{N-f}\sum_i \frac{\left(\delta_i^{\text{model}} - \delta_i^{\text{exp}}\right)^2}{s_i^2}; \tag{4}$$

The summation *i* runs over all *N* measured/predicted NMR shifts, *f* is the number of adjustable model parameters (0 for the direct ranking of models; 1 for the exchange model with optimized populations), $\delta_i^{exp}$ is the experimental chemical shift, $\delta_i^{model}$ is the corresponding predicted shift for a given model, and $s_i^2$ is the nuclide-specific weighting derived by setting $s_i$ to the corresponding root-mean-square error derived from benchmark studies. For the 11 experimental chemical shift measurements here for E(A-A), single-site models ($N$-$f$ = 11) with reduced-$\chi^2$ greater than 1.79 and two-site exchange models ($N$-f = 10) with reduced-$\chi^2$ greater than 1.83 can be ruled out with better than 95% confidence(16). For the 13 experimental chemical shift measurements for E(A-A)(BZI), single-site models ($N$-$f$ = 13) with reduced-$\chi^2$ greater than 1.72 and two-site exchange models ($N$-f = 12) with reduced-$\chi^2$ greater than 1.75 can be ruled out with better than 95% confidence.

Protein Preparation

Tryptophan synthase was prepared by overexpression of *St*TS in *E. coli* BL21 celss as previously described (17, 18). Samples were prepared with the following isotopic labeling schemes for the cofactor and protein components: (1) Protein and cofactor unlabeled/natural abundance isotopomer concentration; (2) Protein $^{15}$N-labeled at lysine ε-nitrogen sites (ε-$^{15}$N-Lys TS); and (3) Protein with the PLP cofactor selectively $^{13}$C enriched at carbon sites C2, C2', and C3 and $^{15}$N enriched at the pyridine ring nitrogen (2,2',3-$^{13}C_3$,$^{15}$N-PLP; TS). Isotopically labeled PLP was prepared as previously detailed (and elaborated upon below) and exchanged into the β-subunit active site as detailed in (17).

Microcrystalline Protein Samples for Solid-State NMR

Microcrystalline samples of TS were prepared by diluting enzyme solution 1:1 with 50 mM $Cs^+$-bicine buffer, pH 7.8, containing 14% PEG-8000 and 3.0 mM spermine (17). Microcrystals were collected and washed with 50 mM $Cs^+$-bicine, pH 7.8, containing 8% PEG-8000, 1.8 mM spermine, and 3 mM N-(4'-trifluoromethoxybenzenesulfonyl)-2-aminoethyl phosphate (F9; a high affinity alpha site ligand and analogue of the natural α-site substrate 3-idole-D-glycerol-3'-phosphate (IGP) (19)). The crystals were packed at 10,000 rpm into a Bruker 4 mm magic-angle spinning (MAS) rotor with an approximate volume of 80 µL; each rotor contained 25-30 mg of protein. To form the α-aminoacrylate intermediate, serine was introduced by direct addition of 5 µL of 1.2 M L-Ser to the packed MAS rotor. Stabilization of the α-aminoacrylate species is enhanced by low temperature (-5 °C), the use of the tight binding α-subunit ligand F9, and the presence of $Cs^+$ (19), which binds to the monovalent cation site in the β-subunit.

X-Ray Crystallography

The X-ray crystal structures of *St*TS E(A-A) and *St*TS E(A-A)(BZI) with $NH_4^+$/$Cs^+$ bound to the monovalent cation site with N-(4'-trifluoromethoxybenzenesulfonyl)-2-aminoethyl phosphate (F9) bound to the α-site were solved at 1.4 Å resolution ($NH_4^+$,F9; PDBID 7MT4), 1.50 Å resolution ($Cs^+$, F9; PDBID 7MT5), and 1.70 Å resolution ($Cs^+$, F9, BZI; PDBID 7MT6) using previously described protocols (4, 20). In brief, crystals were grown at room temperature by sitting drop vapor diffusion by mixing 5.0 µL of the protein solution at 15 mg/mL containing 2.0 mM of the inhibitor (F9) (19) and 5 µL of reservoir solution containing 50 mM Cs-bicine pH 7.8 ($Cs^+$ form)/Na-bicene pH 7.8 ($NH_4^+$ form), 2 mM spermine, and 8-10% V/V PEG8000. Crystals grew to maximum dimensions within two weeks. To prepare the aminoacrylate form, 25 mM L-Serine was mixed in an aliquot of the reservoir buffer (buffer L-Ser). To cryoprotect the *St*TS crystals, PEG400 (10-30%) was mixed with L-Ser buffer, quick-soaked and flash-frozen. The data set was indexed, integrated and scaled using the CCP4 platform (21). The complex model was improved using iterative cycles of manual rebuilding with the program Coot (22), and refinement using REFMAC (23) and PHENIX (24). The data collection and refinement statistics are summarized in Table S3.

Observation of $NH_4^+$ bound in the monovalent cation binding site was unexpected. The $Na^+$ form of E(A-A) produces ammonium pyruvate at an accelerated rate relative to the $Cs^+$ form, and we hypothesize that the $NH_4^+$ displaces the $Na^+$, which binds more weakly than $Cs^+$ (25-27).

NMR Spectroscopy

*$^{13}$C and $^{15}$N Solid-State NMR Spectroscopy:* $^{13}$C and $^{15}$N cross-polarization (CP) magic-angle-spinning (MAS) experiments were performed at 9.4 T (400.37 MHz $^1$H, 100.69 MHz $^{13}$C, 40.57 MHz $^{15}$N) on a Bruker AVIII

spectrometer equipped with a double resonance, 4 mm MAS probe, spinning at MAS rates of 8 kHz; the bearing gas was cooled to -15 °C, giving an effective sample temperature of -10 °C. Cross-polarization was accomplished at a $^{1}H$ spin-lock field of 45 kHz and a $^{13}C/^{15}N$ spin-lock of 54 kHz ($^{13}C$) and 37 kHz ($^{15}N$) (ramped +/- 2 kHz); 85 kHz Spinal64 $^{1}H$ decoupling (28) was used throughout. $^{13}C$ spectra consist of the sum of 16,384 transients acquired with a relaxation delay of 4 s, for a total acquisition time of 18.3 h. $^{13}C$ chemical shifts were referenced indirectly to neat TMS via an external solid-state sample of adamantane with the downfield-shifted peak set to 38.48 ppm (29, 30). $^{15}N$ spectra consist of the sum of 81,920 transients acquired with a relaxation delay of 4 s, for a total acquisition time of 3 d 19 h. $^{15}N$ chemical shifts were referenced indirectly to liq-$NH_3$ (25 °C) via an external solid-state sample of $^{15}NH_4Cl$, in which the resonance frequency was set to 39.27 ppm (29).

The acquisition of solid-state NMR spectra was interleaved with single pulse, low-power decoupling experiments (64 scans $^{13}C$, 256 scans $^{15}N$) reporting predominantly on free ligand and reaction products in solution (mother liquor) (Fig. S4). Acquisition of solid-state NMR spectra for the intermediate was halted before reactant concentrations in solution fell to zero.

*$^{31}P$ Solid-State NMR Spectroscopy:* $^{31}P$ CPMAS experiments on the α-aminoacrylate species were performed at 14.1 T (600.01 MHz $^{1}H$, 242.89 MHz $^{31}P$) on a Bruker NEO spectrometer equipped with an $^{1}H$-X double resonance 4 mm MAS probe, spinning at a MAS rate of 10 kHz. The bearing gas was cooled to -15 °C, giving an effective sample temperature of -5 °C. Cross-polarization was accomplished with a $^{1}H$ spin-lock field of 45 kHz and a $^{31}P$ spin-lock of 47 kHz (ramped +/- 5 kHz); 58 kHz Spinal64 $^{1}H$ decoupling (28) was used during detection. The $^{31}P$ spectra consist of the sum of 8,192 transients acquired with a relaxation delay of 4 s, for a total acquisition time of 9.1 h. $^{31}P$ chemical shifts were indirectly referenced to 85% $H_3PO_4$ (MAS). For comparison to measurements in solution, δ[85% $H_3PO_4$ (capillary)] = δ[85% $H_3PO_4$ (sphere/MAS)] + 0.36 ppm (31).

*$^{15}N$-observe, $^{31}P$-dephased Rotational Echo Double Resonance Experiments:* $^{15}N(^{31}P)$-REDOR(32) experiments were performed at 21.1 T (898.66 MHz $^{1}H$; 91.06 MHz $^{15}N$; 363.78 MHz $^{31}P$) on a Bruker AVANCE 900 spectrometer equipped with an NHMFL low-E, triple resonance 3.2 mm MAS probe(33) (sample volume ~ 40 µl) and spinning at a MAS rate of 8 kHz. The bearing gas was cooled to -15 °C, giving an effective sample temperature of -10 °C. Cross-polarization was accomplished at a $^{1}H$ spin-lock field of 45 kHz, $^{15}N$ spin-lock of 37 kHz (ramped +/- 2 kHz), and a 2 ms contact time; 100 kHz Spinal64 $^{1}H$ decoupling(34) was used throughout. A single 10 µs π pulse was applied to $^{15}N$ at the center of the 25 ms echo period, while a series of 12 µs π pulses at half rotor intervals were applied to $^{31}P$ during the dephasing (S) experiments. The REDOR S and $S_o$ spectra were acquired in an interleaved fashion, and each spectrum consists of the sum of 28,672 transients acquired with a relaxation delay of 2 s, for a total acquisition time of 18.2 h

*$^{15}N$ Chemical Shift Anisotropy Measurements:* Slow spinning $^{15}N$ CPMAS experiments on E(A-A) and E(A-A)(BZI) were performed at 14.1 T (600.01 MHz $^{1}H$, 60.82 MHz $^{15}N$) on a Bruker NEO spectrometer equipped with an $^{1}H$-X double resonance 4 mm MAS probe, spinning at a MAS rate of 2.3 kHz. The bearing gas was cooled to -15 °C, giving an effective sample temperature of -10 °C. Cross-polarization was accomplished with a $^{1}H$ spin-lock field of 45 kHz and a $^{15}N$ spin-lock of 47 kHz (ramped +/- 5 kHz); 83 kHz Spinal64 $^{1}H$ decoupling (28) was used during detection. $^{15}N$ spectra consist of the sum of 81,920 transients acquired with a relaxation delay of 4 s, for a total acquisition time of 3 d 19 h.

*Chemical Shift Tensor Analysis:* Chemical shift tensor principal axis components were determined by a fit of the sideband intensities in the slow MAS spectra using Herzfeld-Berger analysis (35), implemented within Bruker BioSpin's *Topspin 3.6* processing software.

2D $^{13}C$ Correlation Spectroscopy on a CPMAS CryoProbe

2D $^{13}C$-$^{13}C$ CORD (36) were collected at 14.1 T (600.03 MHz $^{1}H$, 150.9 MHz $^{13}C$) on a standard bore Bruker NEO spectrometer equipped with a 3.2 mm CPMAS cryoprobe (Bruker Biosolids CryoProbe™) in which the detection coil and preamplifier operate at cryogenic temperatures to increase sensitivity while the sample remains at lab temperature (37). Experiments were performed at a MAS rate of 13.5 kHz. Cross Polarization was accomplished with a $^{13}C$ spin lock of 55 kHz and $^{1}H$ spin lock of 68 kHz (linearly ramped ±15%). 62.5 kHz $^{13}C$ and 100 kHz $^{1}H$ pulses were used throughout and 100 kHz Spinal64 $^{1}H$ decoupling (28) was used during detection. The CORD mixing time was 50ms. In the indirect dimension, 160 complex points with a dwell of 37.04 μs (spectral width 27 kHz, total acquisition time 5.93 ms) were acquired. In the direct dimension, 1024 complex points with a dwell of 22 μs (spectral width 45.45 kHz, total acquisition time 22.52 ms) were acquired. 16 transients at each $t_1$ point were coadded with a relaxation delay 2 s for a total experiment time of 2.84 hr. The sample temperature was regulated with Bruker variable temperature controller set to -21 °C, giving an effective sample temperature of -10 °C.

$^{15}N$ DNP Experiments

MAS-DNP NMR spectra were collected at 600 MHz (14.1 T) using a Bruker AVIII DNP NMR system (Bruker Biospin) equipped with a Bruker 3.2 mm, $^{1}H/^{13}C/^{15}N$ MAS-DNP probe cooled to ~95 K. Microwaves were delivered using a gyrotron source operating at 395 GHz and output optimized for ~12 W at the probe base (38). Microwaves on/off spectra were collected with a MAS frequency of 10 kHz and a $^{13}C$ CP-echo pulse sequence with a $^{1}H$ CP ramp from 40 to 70 kHz, a $^{13}C$ RF field of 50 kHz, 85 to 100 kHz Spinal64 $^{1}H$ decoupling (28) and a recycle delay of 5s. An AsymPolPOK (39, 40) concentration of ~5 mM was used, resulting in enhancements >60 and buildup times on the order of 3 s.

*Evaluation of MAS-DNP samples by X-band EPR spectroscopy:* CW EPR spectra were collected on a Bruker X-Band EMX Nano benchtop spectrometer (Bruker Biospin, Billerica, MA) to verify polarizing agent concentrations and PA dispersion upon freezing as described in Tran et al. (38). DNP sample rotors were placed in 5 mm quartz sample tubes and placed into the cavity with its associated adaptor. Spectra were collected at room temperature (298 K) and at 100 K by using cold nitrogen gas to cool the cavity.

$^{17}O$ QCT NMR Experiments and the Series Connected Hybrid

$^{17}O$ QCT NMR experiments on E(A-A) and E(A-A)(BZI) were conducted at temperatures between 2 °C and 5 °C at field strengths of 11.7 T, 14.1 T, 16.4 T, 21.1T, and 35.2T. The 35.2T is the Series Connected Hybrid (SCH) magnet at the National High Magnetic Field Laboratory (NHMFL) in Tallahassee (41). All spectra were referenced to the natural abundance $^{17}O$ signal in an external sample of pure water measured at 25°C and assigned the value of 0 ppm. 90° pulse width calibrations were first determined using the natural abundance $^{17}O$ signal of water; the nutation frequencies were 9 kHz, 30 kHz, 46 kHz, 60 kHz, and 60 kHz for field strengths of 11.7 T, 14.1 T, 16.4 T (BZI sample only), 21.1T, and 35.2T respectively. Simultaneous mode quadrature detection was used to acquire $^{17}O$ spectra at all fields measured. A total of 2048 or 4096 points were collected and zero-filled to two times the number of real points, spectral widths ranged from 41 kHz to 100 kHz, relaxation delays were between 5 and 10 ms, and acquisition times were between 10 and 25 ms. A 90°- τ -180°- τ -acquire pulse sequence was used to measure the spectra of all intermediates. A 90°-τ-180°-2τ-180°-2τ-180°-τ -acquire (triple-echo, 3π) was additionally used to measure the spectra of the E(A-A) and E(A-A)(BZI) intermediates at 14.1T and higher to improve observation of bound resonances by suppressing signal from the free substrate and reaction product (42). This is based on the selective excitation of the central transition in $^{17}O$ QCT NMR which leads to a nutation rate that is three times faster for protein-bound substrates than that of the broadband excitation felt by free substrate in solution (43, 44). The power levels for the 180° refocusing pulses were set to those of the 90° pulses for 11.7T, 14,1T, and 16.4T, but were lowered to 10 kHz for 21.1T and 35.2T. In each case the 180° pulse length was set to 1/3 the nominal pulse for water; for 11.7T, 14.1T, and 16.4T, the initial excitation pulse was set to 1/3 the water 90° pulse, while at 21.1T and 35.2T it was set to 1/2 the water 90°. For all experiments, the τ delay times were between 5 and 50 μs and generally kept as short as the instrumentation timing controller could handle. The pulse diagrams and phase cycling lists are provided in our previous publication (42). All spectra were ultimately processed using Bruker's Topspin 3.6 or

4.0 software. Field specific details including hardware are listed below.

11.7 T (500.05 MHz $^1$H, 67.79 MHz $^{17}$O): Varian $^{UNITY}$Inova spectrometer equipped with a 5 mm SW PFG probe. Approximately 350 µL of sample was contained in 5 mm Shigemi restricted volume NMR tubes magnetic susceptibility matched to $D_2O$ (Shigemi Inc.).

14.1 T (600.01 MHz $^1$H, 81.34 MHz $^{17}$O): Bruker Avance I spectrometer equipped with a 5 mm BBO Z-grad probe. Approximately 350 µL of sample was contained in 5 mm Shigemi restricted volume NMR tubes magnetic susceptibility matched to $D_2O$.

16.4 T (699.69 MHz $^1$H, 94.85 MHz $^{17}$O): Bruker Avance I spectrometer equipped with a modified Alderman-Grant coil probe (MAGC) built and tuned to $^{17}$O by C. V. Grant at the Center for NMR Spectroscopy and Imaging of Proteins, University of California, San Diego(45). Approximately 200 µL of sample was contained in 5 mm x 1 cm flat bottom glass NMR tubes.

21.1 T (900 MHz $^1$H, 121.56 MHz $^{17}$O): Bruker Avance III spectrometer equipped with a low-E static double resonance probe with a 5 mm round coil. Both magnet and probe were built in-house at the National Magnetic Field Laboratory in Tallahassee, FL. (46). Approximately 200 µL of sample was contained in 5 mm x 1 cm flat bottom glass NMR tubes.

35.2 T (1600 MHz $^1$H, 203.36 MHz $^{17}$O): Bruker NEO spectrometer equipped with a low-E static double resonance probe containing a 4 mm solenoid coil built in-house at the National Magnetic Field Laboratory in Tallahassee, FL (41). Approximately 75 µL of sample was contained in 4 mm OD cylindrical holder.

The $^{17}$O QCT data are shown in Figs. S2 and S3, accompanied by the data analysis.

<u>Synthesis of $^{15}$N-Benzimidazole:</u>

$^{15}$N-benzimidazole (BZI) was synthesized from $^{15}NH_4OH$ and 1-fluoro-2-nitrobenzene in the following sequence:

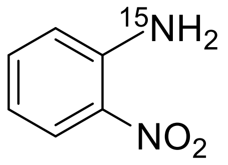


*Synthesis of 2-Nitro-$^{15}$N-aniline:*

To a sealed tube was added 1-fluoro-2-nitrobenzene (105 µL, 1.0 mmol), $^{15}NH_4OH$ solution (303 µL, 1.0 mmol, 3.3 M in water), and Cu(I)Br (14.3 mg, 0.1 mmol). The material was dissolved in 1 mL of bis(2-methoxyethyl) ether, sealed and heated to 120 °C for 16 h. The reaction was then cooled to room temperature, diluted with $CH_2Cl_2$ (5 mL) and the aqueous layer removed. The organic layer was concentrated in vacuo to yield 2-Nitro-$^{15}$N-aniline as a red crystalline solid (57 mg, 41 %). $^1$H NMR (500 MHz, $CDCl_3$) δ 8.12 (d, *J* = 8.6 Hz, 1H), 7.37 (t, *J* = 7.3 Hz, 1H), 6.84 (d, *J* = 8.0 Hz, 1H), 6.71 (t, *J* = 7.8 Hz, 1H), 6.18 (s, 1H), 6.00 (s, 1H). $^{13}$C NMR (126 MHz, $CDCl_3$) δ 144.73, 144.60, 135.66, 132.30, 126.19, 118.85, 116.95.

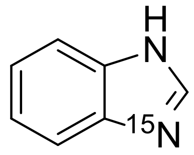


*Synthesis of $^{15}$N-Benzimidazole:*

2-Nitro-$^{15}$N-aniline (30 mg, 0.2 mmol) was added to a two-neck round bottom flask containing Fe shavings (60 mg, 1.07 mmol), triethyl orthoformate (358 µL, 2.15 mmol), $Yb(OTf)_3$ (1 mg, 0.5 mol %) and 1

M AcOH (215μL). The reaction was heated to 75 °C for 24 h. The mixture was then cooled to room temperature, filtered through a celite plug, diluted with 3 mL DI water and extracted with $CH_2Cl_2$ (2 x 4 mL). The resulting solution was concentrated in vacuo to give a yellow residue. The product was further purified through column chromatography in using 1% MeOH/$CH_2Cl_2$ eluent to afford a white solid (10 mg, 42 %). $^1$H NMR (400 MHz, $CDCl_3$) δ 8.67 (s, 1H), 8.18 (d, *J* = 9.6 Hz, 1H), 7.75 – 7.66 (m, 2H), 7.36 – 7.29 (m, 2H). HRMS (ESI) m/z calcd. for $C_7H_6N^{15}N$: 119.0501, found 118.0460 $(M-H)^-$.

Synthesis of Pyridoxal-5'-Phospahte

$^{15}$N-PLP was prepared from $^{15}$N-Ala by forming first the labeled PL and performing the phosphorylation step enzymatically with the ePL Kinase K229Q mutant (47), kindly provided by Martin Safo. The following sequence was followed:

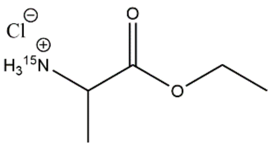


*($^{15}$N)-ethyl 2-aminopropanoate:*

($^{15}$N)-Alanine (750 mg, 6.4 mmol) was dissolved in 20 mL of absolute ethanol inside a round bottom flask and cooled to 0 °C. While stirring, thionyl chloride (4.6 mL, 64.0 mmol) was added dropwise, then the resulting solution was heated to 40 °C for 12 hrs. The reaction was stripped of solvent and volatiles under reduced pressure, allowing the product to spontaneously crystallize inside the flask as (15N)-ethyl-2-aminopropanoate hydrochloride (973 mg, 6.3 mmol, 99% yield). The resulting crystals were used without further purification. Spectroscopic data was consistent with literature. 1H NMR (600 MHz, CDCl3) δ 8.68 (dd, J = 73.4, 4.7 Hz, 3H), 4.30 – 4.16 (m, 3H), 1.72 (dd, J = 7.1, 2.6 Hz, 3H), 1.30 (t, J = 7.1 Hz, 3H).

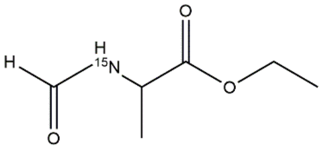


*($^{15}$N)-ethyl 2-formamidopropanoate:*

($^{15}$N)-ethyl-2-aminopropanoate hydrochloride (973 mg, 6.3 mmol) was dissolved in triethyl orthoformate (10 mL, 60 mmol) and refluxed at 140 °C for 2 hrs. The reaction was cooled, then stripped of solvent under reduced pressure, yielding ($^{15}$N)-ethyl 2-formamidopropanoate (890 mg, 6.1 mmol, 97% yield) as a yellow oil. The crude oil was used without further purification. Spectroscopic data was consistent with literature. $^1$H NMR (600 MHz, $CDCl_3$) δ 8.18 (d, J = 16.3 Hz, 1H), 6.27 (dd, J = 91.8, 7.5 Hz, 1H), 4.65 (p, J = 7.2 Hz, 1H), 4.21 (q, J = 7.1 Hz, 2H), 1.44 (dd, J = 7.1, 2.7 Hz, 3H), 1.29 (t, J = 7.1 Hz, 3H).

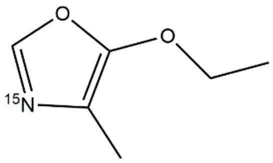


*($^{15}$N)-5-ethoxy-4-methyloxazole*

To a vigorously stirred heterogeneous mixture of phosphorous pentoxide (3.6 g, 12.7 mmol) and magnesium oxide (1.2 g, 30 mmol) in 50 mL of dry dichloromethane, under nitrogen atmosphere, was added a solution of ($^{15}$N)-ethyl 2-formamidopropanoate (600 mg, 4.1 mmol) dissolved in 3 mL of dry dichloromethane. The reaction was refluxed for 24 hrs, then cooled to 0 °C. To the cooled mixture was slowly added 10 mL of water, followed by 50 mL of saturated $NaHCO_3$. The biphasic mixture was filtered through celite, then transferred to a separatory funnel, and the organic phase separated. The aqueous phase was further extracted with dichloromethane (3 x 15 mL). The combined organic layers were dried over $Na_2SO_4$, filtered through cotton, and the solvent removed under reduced pressure to yield ($^{15}N$)-5-ethoxy-4-methyloxazole (260 mg, 2.0 mmol, 49% yield) as a yellow oil. The crude oil was used without further purification. Spectroscopic data was consistent with literature. $^1H$ NMR (500 MHz, $CDCl_3$) δ 7.37 (d, J = 12.6 Hz, 1H), 4.14 (q, J = 7.1 Hz, 2H), 2.04 (d, J = 2.2 Hz, 3H), 1.35 (t, J = 7.1 Hz, 3H).

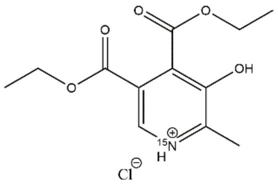


*($^{15}N$)-diethyl 5-hydroxy-6-methylpyridine-3,4-dicarboxylate*

($^{15}N$)-5-ethoxy-4-methyloxazole (260 mg, 2.0 mmol) and diethyl maleate (345 mg, 2.0 mmol) were added to a small pear-shaped flask, and the neat mixture was heated at 75° C for 18 hrs. The reaction was cooled to room temperature, diluted with 5 mL of isopropanol, acidified with HCl (4M in dioxane), then diluted with 100 mL of diethyl ether and placed in the freezer at -20 °C to promote crystallization. The resulting precipitate was filtered and washed with diethyl ether to yield ($^{15}N$)-4,5-bis(ethoxycarbonyl)-3-hydroxy-2-methylpyridin-1-ium chloride (266 mg, 0.9 mmol , 46% yield) as a tan powder. Spectroscopic data was consistent with literature. $^1H$ NMR (500 MHz, DMSO) δ 10.03 (s, 1H), 8.50 (s, 1H), 4.30 (q, J = 7.1 Hz, 2H), 4.28 (q, J = 7.1 Hz, 2H), 2.49 (s, 3H), 1.28 (t, J = 7.1 Hz, 3H), 1.27 (t, J = 7.1 Hz, 3H).

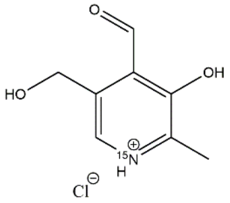


*($^{15}N$)-pyridoxal*

To a stirred solution of ($^{15}N$)-4,5-bis(ethoxycarbonyl)-3-hydroxy-2-methylpyridin-1-ium chloride (100 mg, 0.3 mmol) in 20 mL of dry tetrahydrofuran, under nitrogen atmosphere at 0° C, was added 3.0 mL of $LiAlH_4$ (1M in THF) and the resulting mixture was then refluxed for 24 hrs. The mixture was cooled, then carefully quenched with 25 mL of water, and filtered through celite. The filtrate was washed with dichloromethane (3 x 10 mL), and the aqueous phase was transferred to a round bottom flask along with manganese dioxide (42 mg, 0.5 mmol). The mixture was acidified with aqueous HCl, and allowed to stir at room temperature for 4 hours. The mixture was then filtered through celite, yielding an aqueous solution of crude ($^{15}N$)-pyridoxal (21 mL, 15.7 mM) and the solution was used without further purification. UV-Vis showed peaks at 320 and 388 nm.

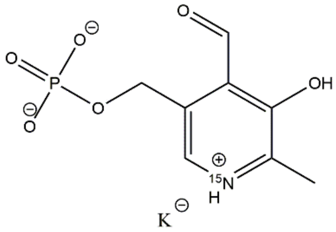


$^{15}N$ pyridoxal-5'-phosphate:

A 5ml solution composed of 32 μM of K229Q PL Kinase, 32 mM ATP, 50 mM KCl, 200 mM HEPES•K pH 7.5,

0.2 mM MgCl, and 5 mM ($^{15}N$)-pyridoxal (15.7 mM) was stirred and the reaction run overnight at room temperature. A 1mL aliquot of the solution was monitored by UV-Vis at 388 nm, showing the buildup of PLP. The crude solution was used without further purification for exchange into the apoenzyme as previously described (17). Spectra for the holoenzyme were consistent with literature.

The K229Q mutant ePL kinase was expressed and purified according to (48).

Scheme S1: α-aminoacrylate intermediate candidate structures

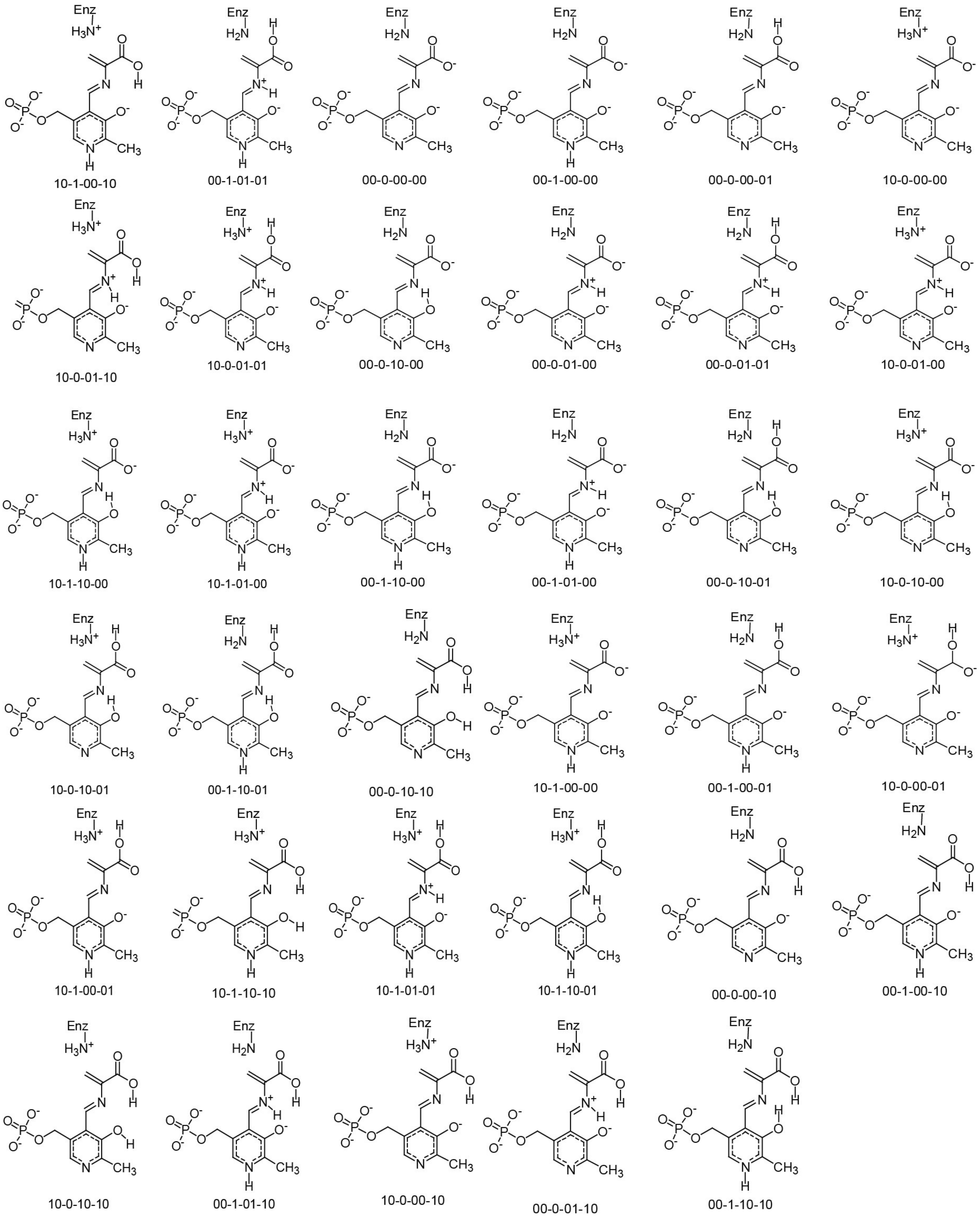

Scheme S2: E(A-A)(BZI) candidate structure

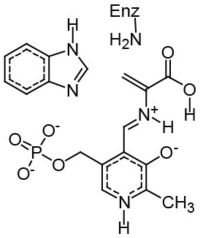
00-1-01-10

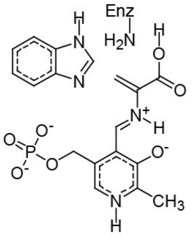
00-1-01-01

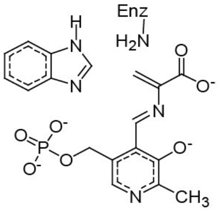
00-0-00-00

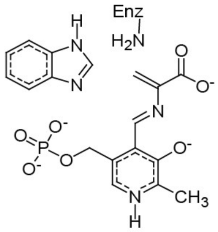
00-1-00-00

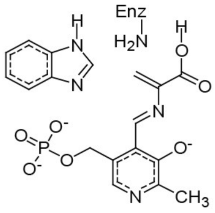
00-0-00-01

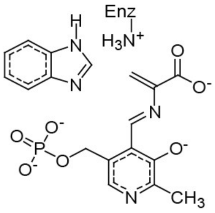
10-0-00-00

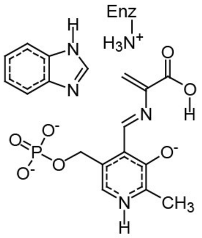
10-1-00-10

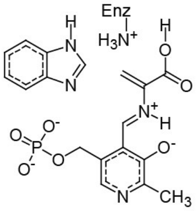
10-0-01-01

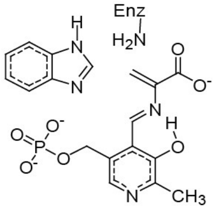
00-0-10-00

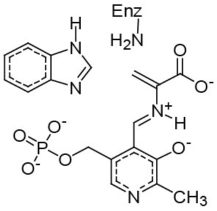
00-0-01-00

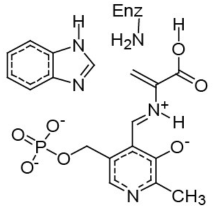
00-0-01-01

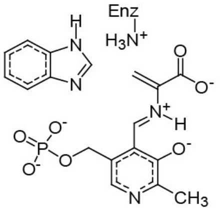
10-0-01-00

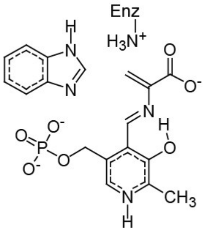
10-1-10-00

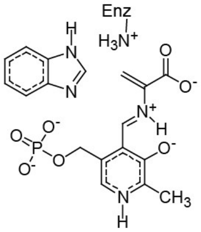
10-1-01-00

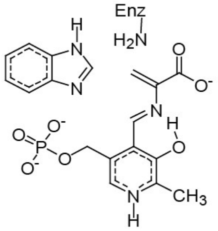
00-1-10-00

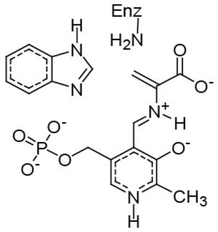
00-1-01-00

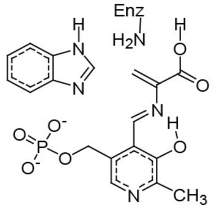
00-0-10-01

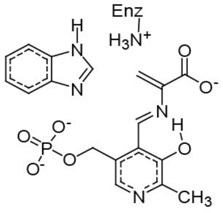
10-0-10-00

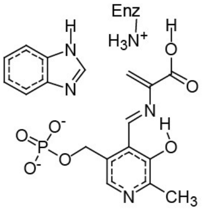
10-0-10-01

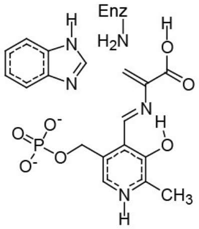
00-1-10-01

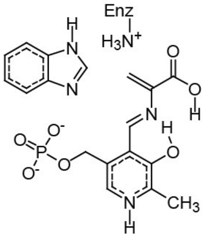
10-1-10-10

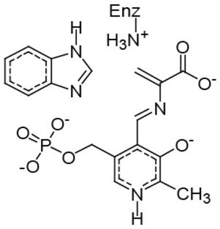
10-1-00-00

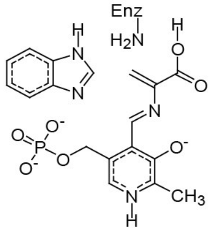
00-1-00-01

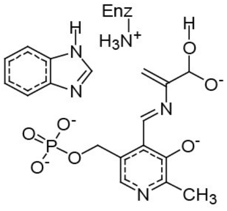
10-0-00-01

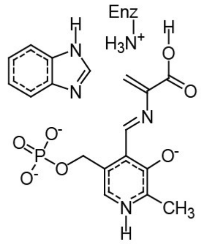
10-1-00-01

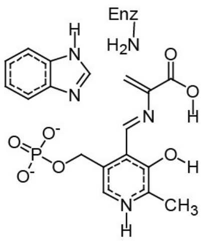
00-1-10-10

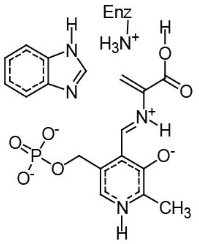
10-1-01-01

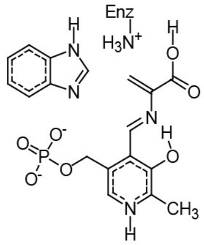
10-1-10-01

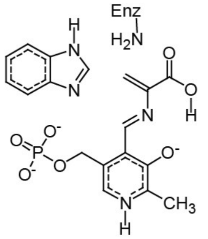
00-1-00-10

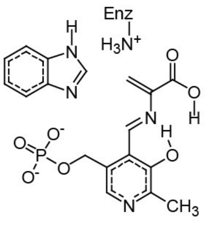
10-0-10-10

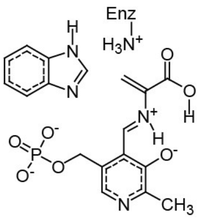
10-0-01-10

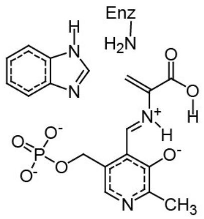
00-0-01-10

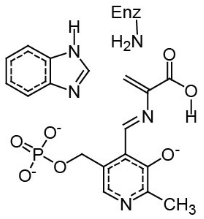
00-0-00-10

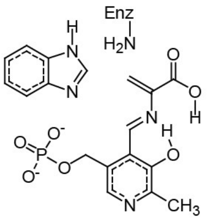
00-0-10-10

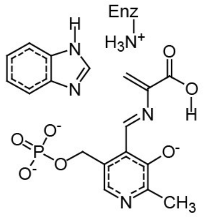
10-0-00-10

Table S1: Key first-principles chemical shifts (ppm) for the E(A-A) intermediate candidates

| | PLP | | | | L-Serine | | | | | | βK87 | |
|---|---|---|---|---|---|---|---|---|---|---|---|---|
| Structure | N1 | C2 | C2' | C3 | SB N | Cα | C' | Cβ | O1 | O2 | Nζ | Red χ2 |
| 00-0-10-00 | 302.9 | 148.7 | 18.8 | 154.9 | 301.4 | 144.4 | 171.2 | 123.3 | 262.1 | 302.4 | 20.5 | 3.1 |
| 10-0-10-00 | 313.5 | 152.5 | 19.4 | 155.3 | 300.9 | 145.8 | 169.6 | 120.4 | 264.1 | 307.5 | 36.0 | 4.1 |
| 00-0-00-00 | 278.9 | 157.7 | 20.3 | 166.4 | 324.3 | 154.5 | 173.6 | 113.8 | 256.2 | 314.6 | 21.7 | 18.8 |
| 10-0-00-00 | 284.4 | 161.7 | 20.9 | 167.1 | 318.0 | 156.6 | 172.4 | 108.6 | 259.1 | 319.4 | 37.8 | 25.2 |
| 00-0-00-01 | 286.4 | 159.0 | 20.4 | 167.4 | 293.6 | 146.7 | 170.6 | 123.5 | 167.3 | 340.2 | 21.2 | 25.2 |
| 10-0-10-01 | 325.0 | 153.4 | 19.2 | 155.1 | 276.0 | 137.5 | 168.9 | 129.7 | 173.0 | 341.2 | 36.7 | 28.9 |
| 00-0-10-01 | 313.4 | 149.5 | 18.6 | 154.5 | 277.7 | 136.0 | 169.8 | 132.7 | 167.9 | 331.9 | 20.4 | 29.6 |
| 10-0-00-01 | 291.4 | 163.0 | 20.8 | 169.0 | 284.8 | 148.2 | 170.5 | 115.7 | 170.1 | 350.7 | 38.0 | 30.7 |
| 10-1-10-00 | 217.8 | 146.1 | 12.9 | 156.6 | 315.4 | 144.0 | 168.4 | 125.5 | 268.0 | 304.0 | 35.8 | 40.5 |
| 10-0-10-10 | 332.6 | 152.7 | 18.6 | 152.4 | 258.2 | 134.1 | 166.6 | 134.6 | 150.4 | 313.4 | 36.9 | 47.9 |
| 00-0-10-10 | 319.5 | 149.1 | 18.2 | 153.5 | 258.7 | 130.7 | 168.4 | 137.3 | 150.9 | 306.2 | 20.5 | 49.2 |
| 00-1-10-00 | 209.6 | 141.9 | 12.3 | 156.1 | 311.8 | 142.2 | 170.1 | 127.3 | 266.7 | 298.8 | 19.9 | 49.6 |
| 00-1-00-10 | 209.6 | 141.9 | 12.2 | 156.1 | 311.7 | 142.2 | 170.2 | 127.3 | 266.7 | 298.8 | 19.9 | 49.6 |
| 10-1-10-01 | 216.1 | 145.3 | 12.8 | 155.7 | 282.0 | 135.4 | 168.5 | 128.4 | 173.5 | 334.6 | 36.5 | 58.7 |
| 10-1-00-01 | 208.8 | 154.3 | 18.1 | 165.7 | 314.9 | 147.2 | 169.4 | 122.2 | 172.4 | 351.8 | 37.9 | 64.7 |
| 00-1-10-01 | 216.3 | 143.2 | 12.4 | 155.9 | 289.2 | 133.8 | 169.4 | 135.8 | 170.2 | 329.6 | 20.1 | 68.6 |
| 00-1-00-01 | 204.7 | 149.5 | 17.6 | 164.5 | 322.9 | 144.6 | 169.8 | 127.7 | 169.6 | 341.3 | 21.3 | 70.6 |
| 10-1-00-10 | 212.8 | 155.7 | 18.1 | 168.7 | 257.4 | 136.0 | 171.9 | 122.3 | 181.1 | 290.4 | 37.3 | 71.6 |
| 10-1-00-00 | 204.1 | 151.9 | 18.1 | 164.3 | 350.1 | 155.6 | 170.7 | 112.8 | 263.9 | 318.4 | 37.5 | 71.9 |
| 00-1-00-00 | 200.0 | 147.3 | 17.6 | 163.9 | 354.2 | 152.9 | 172.1 | 117.4 | 261.7 | 313.2 | 21.8 | 74.3 |
| 10-1-10-10 | 215.6 | 143.5 | 12.2 | 152.4 | 257.8 | 132.8 | 166.3 | 130.7 | 146.0 | 310.9 | 36.5 | 76.8 |
| 00-1-10-10 | 219.7 | 142.9 | 15.1 | 155.6 | 270.6 | 128.5 | 167.3 | 140.7 | 148.4 | 316.0 | 20.2 | 86.5 |
| 00-0-00-10 | 303.3 | 159.1 | 19.8 | 173.2 | 153.6 | 133.1 | 168.7 | 121.3 | 261.5 | 294.3 | 21.1 | 106.1 |
| 00-0-01-00 | 303.3 | 159.1 | 19.8 | 173.2 | 153.6 | 133.1 | 168.7 | 121.3 | 261.4 | 294.3 | 21.1 | 106.1 |
| 10-0-01-00 | 309.4 | 162.7 | 20.3 | 174.4 | 148.0 | 135.7 | 166.9 | 116.5 | 264.0 | 297.3 | 37.2 | 117.4 |
| 10-0-00-10 | 309.3 | 162.7 | 20.3 | 174.4 | 148.0 | 135.7 | 166.9 | 116.5 | 263.9 | 297.3 | 37.2 | 117.4 |
| 00-1-01-00 | 213.6 | 151.7 | 17.0 | 170.7 | 169.1 | 131.2 | 167.4 | 127.0 | 266.2 | 291.6 | 21.7 | 120.8 |
| 10-1-01-00 | 220.1 | 156.5 | 17.7 | 172.1 | 163.5 | 134.3 | 165.4 | 123.2 | 268.2 | 294.7 | 37.0 | 121.4 |
| 00-0-01-01 | 313.6 | 159.2 | 19.8 | 174.0 | 141.9 | 126.4 | 167.8 | 129.5 | 168.7 | 327.6 | 21.4 | 152.3 |
| 10-0-01-01 | 319.8 | 162.9 | 20.3 | 175.7 | 135.7 | 129.5 | 166.7 | 124.6 | 174.8 | 334.7 | 37.7 | 160.4 |
| 00-1-01-01 | 219.1 | 152.6 | 16.5 | 171.8 | 156.7 | 124.6 | 167.1 | 134.7 | 171.7 | 326.3 | 22.3 | 163.7 |
| 00-0-01-10 | 322.7 | 156.6 | 20.3 | 170.5 | 150.0 | 122.0 | 166.0 | 140.6 | 152.1 | 316.3 | 22.1 | 164.6 |
| 10-1-01-01 | 216.1 | 154.0 | 16.9 | 171.9 | 145.6 | 128.0 | 166.2 | 123.9 | 176.0 | 329.7 | 37.5 | 167.1 |
| 10-0-01-10 | 332.2 | 160.1 | 20.8 | 171.9 | 143.7 | 126.2 | 163.7 | 137.0 | 151.7 | 328.8 | 37.3 | 169.8 |
| 00-1-01-10 | 221.8 | 150.1 | 14.0 | 169.2 | 162.1 | 119.6 | 164.5 | 143.9 | 149.1 | 324.4 | 23.3 | 185.3 |
| Structures with 2 waters (center water removed) | | | | | | | | | | | | |
| 00-0-01-00-2W | 301.5 | 158.9 | 20.1 | 173.0 | 153.0 | 135.8 | 169.0 | 120.9 | 264.0 | 293.9 | 22.3 | 103.86 |
| 00-0-10-00-2W | 301.4 | 148.4 | 19.1 | 154.0 | 305.9 | 148.5 | 171.4 | 122.4 | 264.9 | 301.7 | 21.3 | 4.1 |
| Expt | 297.6 | 151.2 | 17.5 | 158.1 | 286.7 | 145.6 | 170.9 | 118.8 | 292.0 | 258.0 | 24.1 | |

Table S2: Key first-principles chemical shifts (ppm) for the E(A-A)(BZI) complex candidate structures

| Structure | PLP | | | | L-Serine | | | | | | BZI | | βK87 | |
|---|---|---|---|---|---|---|---|---|---|---|---|---|---|---|
| | N1 | C2 | C2' | C3 | SB N | Cα | C' | Cβ | O1 | O2 | N3 | N1 | Nζ | Red χ2 |
| 10-0-10-00 | 310.2 | 152.1 | 19.4 | 155.2 | 307.6 | 147.2 | 168.9 | 120.6 | 261.4 | 301.4 | 231.4 | 167.5 | 35.6 | 2.1 |
| 00-0-10-00 | 303.9 | 149.0 | 18.7 | 154.7 | 307.3 | 145.5 | 169.9 | 121.8 | 258.7 | 298.7 | 253.6 | 162.4 | 21.2 | 6.1 |
| 10-0-00-00 | 284.0 | 161.8 | 20.9 | 166.7 | 326.8 | 156.2 | 171.6 | 109.5 | 258.4 | 311.0 | 236.8 | 163.5 | 37.1 | 19.5 |
| 00-0-00-00 | 281.3 | 158.5 | 20.3 | 166.8 | 324.7 | 155.3 | 172.4 | 111.3 | 255.3 | 308.5 | 259.2 | 158.9 | 22.6 | 20.7 |
| 10-0-10-01 | 321.8 | 153.0 | 19.3 | 154.9 | 281.7 | 138.2 | 168.0 | 130.4 | 166.5 | 337.5 | 232.4 | 170.6 | 36.1 | 24.4 |
| 10-0-00-01 | 291.1 | 163.1 | 21.1 | 168.5 | 291.9 | 147.9 | 169.7 | 117.1 | 166.4 | 345.1 | 237.4 | 165.0 | 37.3 | 24.7 |
| 00-0-00-01 | 288.7 | 159.8 | 20.5 | 168.1 | 293.2 | 146.7 | 169.7 | 121.0 | 164.3 | 338.1 | 259.4 | 160.5 | 24.3 | 26.5 |
| 00-0-10-01 | 313.6 | 149.7 | 18.5 | 154.2 | 284.2 | 133.9 | 168.1 | 131.5 | 162.9 | 326.5 | 255.0 | 166.5 | 23.9 | 30.6 |
| 10-0-10-10 | 329.3 | 152.8 | 19.2 | 153.5 | 261.5 | 135.4 | 166.9 | 136.3 | 147.6 | 303.0 | 230.7 | 172.9 | 36.3 | 39.6 |
| 10-1-10-00 | 196.6 | 146.6 | 13.2 | 156.9 | 330.3 | 146.3 | 167.5 | 127.5 | 261.0 | 301.4 | 228.7 | 171.2 | 35.1 | 40.9 |
| 00-1-10-00 | 192.7 | 142.7 | 12.4 | 156.2 | 329.1 | 143.3 | 168.8 | 127.4 | 258.7 | 297.9 | 252.8 | 165.8 | 21.6 | 51.2 |
| 00-1-00-10 | 208.2 | 141.9 | 12.4 | 155.8 | 321.3 | 142.8 | 168.9 | 126.3 | 262.6 | 295.5 | 251.9 | 164.9 | 21.4 | 51.8 |
| 10-1-00-10 | 209.2 | 154.5 | 18.2 | 167.6 | 278.7 | 140.5 | 167.7 | 120.1 | 160.2 | 326.9 | 230.0 | 168.5 | 35.6 | 56.7 |
| 10-1-10-01 | 216.9 | 145.6 | 12.8 | 156.1 | 289.9 | 135.7 | 167.5 | 132.5 | 167.4 | 332.3 | 230.9 | 172.6 | 35.8 | 58.2 |
| 10-1-00-01 | 206.6 | 154.3 | 18.1 | 165.1 | 322.9 | 146.2 | 168.3 | 122.8 | 167.9 | 346.6 | 233.6 | 167.3 | 36.5 | 60.6 |
| 10-1-00-00 | 190.3 | 152.4 | 13.5 | 163.2 | 363.8 | 155.2 | 170.0 | 114.7 | 258.3 | 311.5 | 233.0 | 166.3 | 35.9 | 66.2 |
| 00-1-00-01 | 203.9 | 150.5 | 17.5 | 165.0 | 322.2 | 144.0 | 168.6 | 126.1 | 166.3 | 338.7 | 256.9 | 162.7 | 24.0 | 67.4 |
| 00-1-00-00 | 188.5 | 148.0 | 12.6 | 163.4 | 360.1 | 153.7 | 170.8 | 116.4 | 256.2 | 308.6 | 256.1 | 161.2 | 22.0 | 69.9 |
| 10-1-10-10 | 216.4 | 143.8 | 15.8 | 154.9 | 267.7 | 135.7 | 163.0 | 134.5 | 338.0 | 138.8 | 228.9 | 171.6 | 35.5 | 73.6 |
| 00-0-01-00 | 304.5 | 159.6 | 20.1 | 173.4 | 155.7 | 134.4 | 167.6 | 121.0 | 257.5 | 290.4 | 252.8 | 163.3 | 21.5 | 95.7 |
| 00-0-00-10 | 304.5 | 159.6 | 20.0 | 173.4 | 155.7 | 134.4 | 167.6 | 121.0 | 257.3 | 290.3 | 252.8 | 163.3 | 21.5 | 95.7 |
| 10-0-01-00 | 307.3 | 162.7 | 20.7 | 174.0 | 152.0 | 136.3 | 166.5 | 118.2 | 260.5 | 291.8 | 231.7 | 167.9 | 36.4 | 97.8 |
| 10-0-00-10 | 307.3 | 162.7 | 20.7 | 174.0 | 152.0 | 136.3 | 166.5 | 118.2 | 260.4 | 291.7 | 231.8 | 168.0 | 36.4 | 97.9 |
| 10-1-01-00 | 200.1 | 157.8 | 14.2 | 171.0 | 167.1 | 134.9 | 164.9 | 125.2 | 258.8 | 289.6 | 229.4 | 171.4 | 36.0 | 108.0 |
| 00-1-01-00 | 212.6 | 152.0 | 17.0 | 170.6 | 172.5 | 131.4 | 166.3 | 127.4 | 262.2 | 287.4 | 252.0 | 166.9 | 22.1 | 112.3 |
| 10-0-01-01 | 318.0 | 162.8 | 20.8 | 175.3 | 139.3 | 129.1 | 166.1 | 126.7 | 168.0 | 332.0 | 232.4 | 170.6 | 36.8 | 138.6 |
| 00-0-01-01 | 314.3 | 159.5 | 20.1 | 173.9 | 144.1 | 124.3 | 166.1 | 129.3 | 162.8 | 322.3 | 254.5 | 167.7 | 24.4 | 140.2 |
| 10-0-01-10 | 329.6 | 159.7 | 21.3 | 172.1 | 147.0 | 126.4 | 164.4 | 141.2 | 149.2 | 316.1 | 230.4 | 173.5 | 36.5 | 148.6 |
| 10-1-01-01 | 216.6 | 154.8 | 17.0 | 172.0 | 150.5 | 126.9 | 165.2 | 128.9 | 169.3 | 326.9 | 230.4 | 171.7 | 36.4 | 151.0 |
| *00-0-01-10 | - | - | - | - | - | - | - | - | - | - | - | - | - | - |
| *00-0-10-10 | - | - | - | - | - | - | - | - | - | - | - | - | - | - |
| *00-1-01-01 | - | - | - | - | - | - | - | - | - | - | - | - | - | - |
| *00-1-01-10 | - | - | - | - | - | - | - | - | - | - | - | - | - | - |
| *00-1-10-01 | - | - | - | - | - | - | - | - | - | - | - | - | - | - |
| *00-1-10-10 | - | - | - | - | - | - | - | - | - | - | - | - | - | - |
| Expt | 302.4 | 153.1 | 18.2 | 158 | 292.3 | 146 | 169.8 | 118.7 | 258 | 287 | 227.8 | 165.5 | 35.58 | |

* Unstable structures during geometry optimization: BZI attached to cofactor

Table S3: E(A-A) and E(A-A)(BZI) Schiff base nitrogen experimental and first-principles chemical shift tensor components (ppm) for the phenolic (phen) and protonated Schiff base (PSB) models and their two-site exchange model with the following populations: E(AA) 89.3% Phen (00-0-10-00),10.7% PSB (00-0-01-00); E(AA)(BZI) 89.4% Phen (10-0-10-00), 10.6% PSB (10-0-01-00)

| Site | comp | C-Phen | C-PSB | two-site | expt |
|---|---|---|---|---|---|
| AA | $\delta_{11}$ | 560.6 | 281.0 | 523.7 | 526.5 |
| | $\delta_{22}$ | 331.4 | 154.4 | 319.0 | 306.1 |
| | $\delta_{33}$ | 11.9 | 24.4 | 14.3 | 31.5 |
| | Red-$\chi^2$ | 12.23 | 435.8 | 2.41 | - |
| BZI | $\delta_{11}$ | 577.2 | 279.0 | 540.3 | 545.8 |
| | $\delta_{22}$ | 334.0 | 151.6 | 321.9 | 328.4 |
| | $\delta_{33}$ | 11.5 | 278.4 | 13.9 | 7.9 |
| | Red-$\chi^2$ | 6.3 | 541.2 | 0.51 | - |

**$^{17}O$ QCT NMR Spectroscopy:**

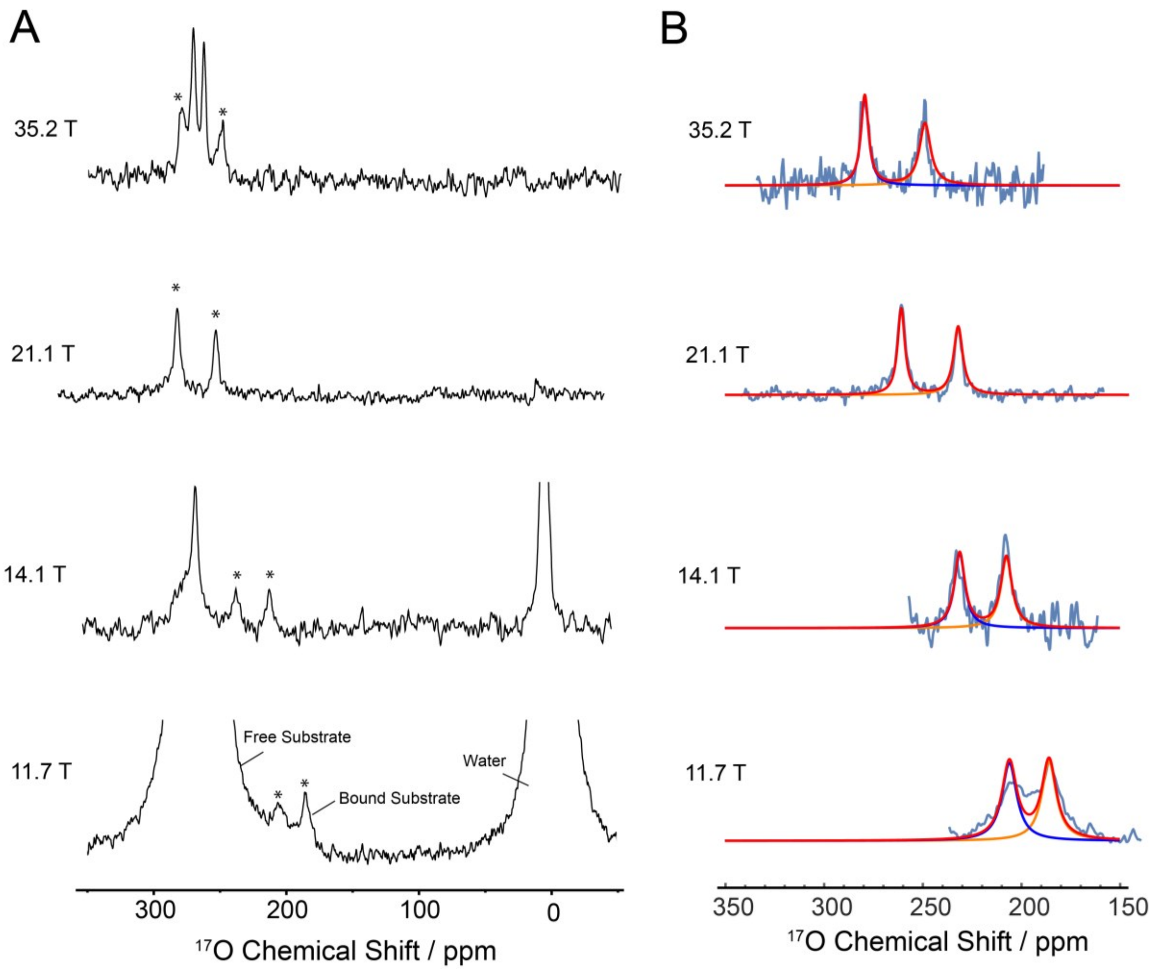


Fig. S2: $^{17}O$ QCT NMR spectra of the E(A-A) intermediate from 11.7 – 35.2 T. At 11.7 T the spectrum was acquired with a single echo before detection and displays considerable intensity from the natural abundance $H_2{}^{17}O$ water and free $^{17}O_2$-L-Ser substrate in solution. At 14.1T and higher, the spectra were acquired using the triple echo sequence to greatly suppress the free substrate and water. (B) Traces of the best-fit component lineshapes O1 (blue), O2 (orange), and their sum O1 + O2 (red), overlaid on experimental spectra at the magnetic field strengths listed. 100 Hz of line broadening was applied to all spectra shown in the figure.

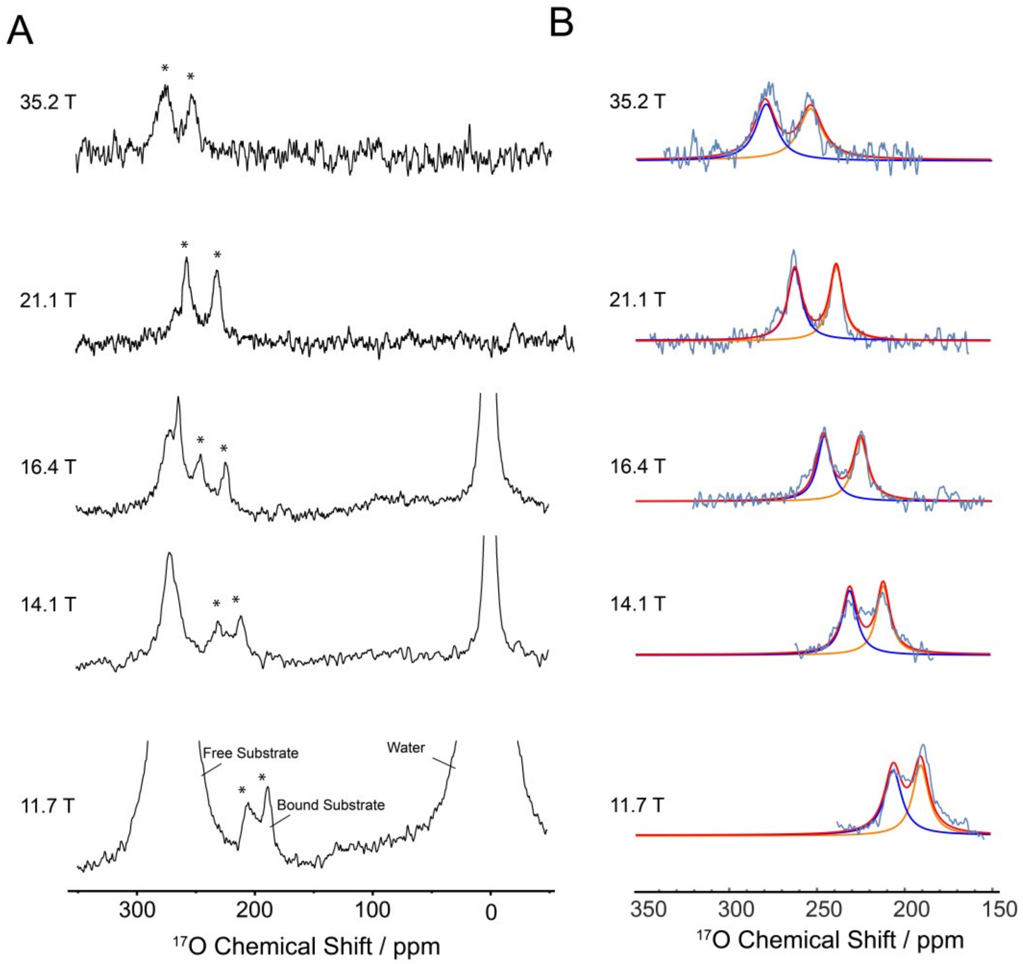


Fig. S3. $^{17}O$ QCT NMR spectra of the E(A-A)(BZI) intermediate from 11.7 – 35.2 T. At 11.7 T the spectrum was acquired with a single echo before detection and displays considerable intensity from the natural abundance $H_2{}^{17}O$ water and free $^{17}O_2$-L-Ser substrate in solution. At 14.1T and higher, the spectra were acquired using the triple echo sequence to greatly suppress the free substrate and water. (B) Traces of the best-fit component lineshapes O1 (blue), O2 (orange), and their sum O1 + O2 (red), overlaid on experimental spectra at the magnetic field strengths listed. 100 Hz of line broadening was applied to all spectra shown in the figure.

***Global Spectral Fitting***:

The precise frequency of the central transition is perturbed from the isotropic value by the second-order quadrupolar interaction, but by measuring the spectrum as a function of field, the isotropic shift and anisotropic quadrupole and chemical shift anisotropy product parameters can be determined. Data fitting to extract the isotropic chemical shifts ($\delta_{iso}$ (ppm)), quadrupole product parameter ($P_Q$ (MHz)), shielding anisotropy product parameter ($P_{SA}$ (ppm)), and rotational correlation time, $\tau_c$, from $^{17}O$ QCT NMR spectra made use of the theory and equations from Wu and coworkers (49, 50), which is briefly summarized here.

In the limit of slow isotropic motion the observed chemical shift, $\delta_{obs}$, appears at a value lower than the $\delta_{iso}$ as a result of 2nd order quadrupolar shift.(51) The 2nd order shift is not observed for $\omega_0\tau_c << 1$, the fast motion limit applicable to small molecules (such as water or the free substrate) having rotational correlation times on the picosecond timescale. This shift is often termed the dynamic frequency shift, $\Delta\delta_d$ and is given by equation (1).(44) $P_Q$, the quadrupole product parameter, contains the nuclear quadrupolar coupling constant, $C_Q$, and asymmetry parameter, $\eta_Q$.(44)

$$\Delta\delta_d(ppm) = \delta_{obs} - \delta_{iso} = -6000\left(\frac{P_Q}{\nu_0}\right)^2 \quad (1)$$

$$P_Q(MHz) = C_Q\left(1+\frac{\eta_Q^2}{3}\right)^{1/2},\ 0 \le \eta_Q \le 1 \quad (2)$$

Measurements of the observed chemical shift at multiple fields can be used to extract $\delta_{iso}$ and $P_Q$ in a straightforward manner given the linear relationship of $\delta_{obs}$ with $\nu_0^{-2}$.

The shielding anisotropy product parameter, $P_{SA}$, and correlation time, $\tau_c$, may be extracted from the field-dependent line shape of the QCT signals according to equation (3). This equation derives from the transverse relaxation properties of the CT in the limit of slow isotropic motion and is specific to spin-5/2 nuclei. The $P_{SA}$ is composed of $\Delta\sigma$ (ppm), the nuclear magnetic shielding anisotropy, and asymmetry parameter, $\eta_{SA}$, as indicated in equation (4). β is the angle between the principal axis systems of the chemical shift and quadrupolar coupling tensors under the assumption of axial symmetry; here we take β=π/2 (49), which is supported by modeling of the tensors using first principles methods.

$$\Delta\nu_{\frac{1}{2}}(Hz) = \frac{R_{obs}^2}{\pi} = 9.6\times10^{-4}\frac{P_Q^4}{\nu_0^2}\tau_c + 7.2\times10^{-3}\left(\frac{P_Q}{\nu_0}\right)^2\frac{1}{\tau_c} + 1.1(P_{SA}\nu_0)^2\tau_c + 0.029\,P_Q^2P_{SA}\tau_c(3cos^2\beta - 1) \quad (3)$$

$$P_{SA}(ppm) = \Delta\sigma\left(1+\frac{\eta_{SA}^2}{3}\right)^{1/2},0 \le \eta_{SA} \le 1 \quad (4)$$

The simultaneous fitting of spectra at multiple magnetic field strengths to equations (1)-(4) allows the extraction of $\delta_{iso}$, $P_Q$, $P_{SA}$, and $\tau_c$. In practice, this gives a Lorentzian line shape for each peak that is described by equation (5) (written to yield units of ppm in the chemical shift dimension). The intensity, $I(\nu_0)$, is not constrained and in practice is given as many free parameters as the number of different field strengths measured. As an example, equation (6) contains a cubic, four free-parameter equation for the intensity of intermediates measured at four different magnetic field strengths as was accomplished for E(A-A).

$$L(I,\nu_0,\delta_{obs},\Delta\nu_{1/2}) = \frac{\nu_0 I(\nu_0)}{\Delta\nu_{1/2}\left(1+4\left(\frac{\nu_0(\delta-\delta_{obs})}{\Delta\nu_{1/2}}\right)^2\right)} \quad (5)$$

$$I(\nu_0) = a + b\nu_0 + c\nu_0^2 + d\nu_0^3 \qquad \text{(4 different field measurements)} \quad (6)$$

Substitution of equations (1)-(4) and (6) into (5) gives an expression for the field-dependent line shape that we used to fit the spectra and extract $\delta_{iso}$, $P_Q$, $P_{SA}$, and $\tau_c$.

To fit the data, the real portion of each spectrum was exported as a JCAMP 6.0 file using Bruker's TopSpin program. The spectra used in the fitting procedure were zero-filled only. No other post acquisition processing was performed although spectra displayed in figures do have line broadening applied, as indicated, for clarity. The JCAMP files were imported into Mathematica and each spectrum was scaled to the same RMSD noise floor. The spectra were then subjected to the global, simultaneous, non-linear least-squares regression to find the best fit parameters satisfying equation (5). $\tau_c$ of 232 ns was used as a fixed parameter in the final fitting of all spectra. This value was first obtained as a result of the simultaneous fitting of the $E(Q_3)_{2AP}$ spectra at 5 field strengths and is consistent with the value expected for the $\tau_c$ of a 143 kDa protein in the 2-5 °C temperature range at which the experiments were conducted.

At each field, the free substrate/product signals were fit and subtracted from the spectrum prior to global fitting of the bound signals. For the SCH E(A-A)(BZI) data only, a reference signal for the free substrate/product was first subtracted. These spectra are shown in Figs. S2 and S3. The best-fit component line shapes of the enzyme-bound $^{17}$O QCT signals are shown overlaid with the experimental spectra (after background peak subtraction) adjacent to the data. The best-fit parameters are tabulated in Table S1.

**Table S4:** $^{17}$O NMR parameters extracted from QCT NMR

| | $\delta_{iso}$/ppm | $P_Q$/MHz | $P_{SA}$/ppm |
|---|---|---|---|
| E(A-A) | | | |
| O1 | 256.6 ± 0.1 | 7.40 ± 0.01 | 341 ± 3 |
| O2 | 288.6 ± 0.1 | 8.00 ± 0.01 | 279 ± 3 |
| E(A-A)(BZI) | | | |
| O1 | 257.8 ± 0.1 | 7.23 ± 0.01 | 505 ± 3 |
| O2 | 285.8 ± 0.1 | 7.89 ± 0.01 | 492 ± 4 |

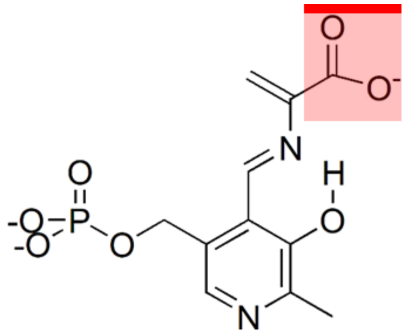

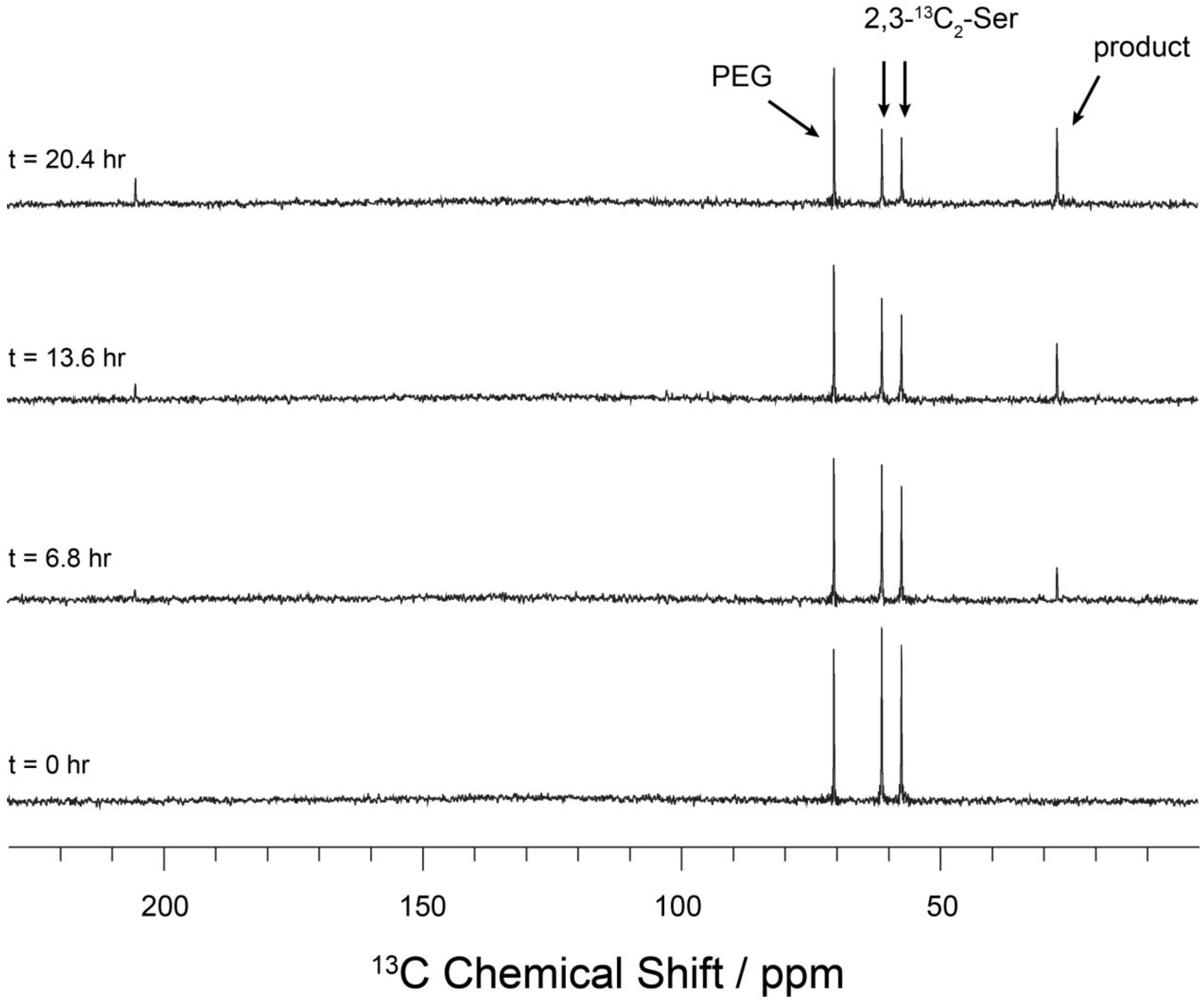


Fig. S4. Direct excitation $^{13}$C solution-state NMR with low power $^{1}$H decoupling spectra of the reaction of tryptophan synthase microcrystals with 10 mM L-[2, 3-$^{13}$C]Ser corresponding to the data shown in Fig. 3. The time course shows the disappearance of serine and appearance of product signals in the mother liquor. Data collected at -10 °C on a 9.4 T Bruker DSX spectrometer equipped with a 4 mm CPMAS probe spinning at a MAS rate of 8 kHz.

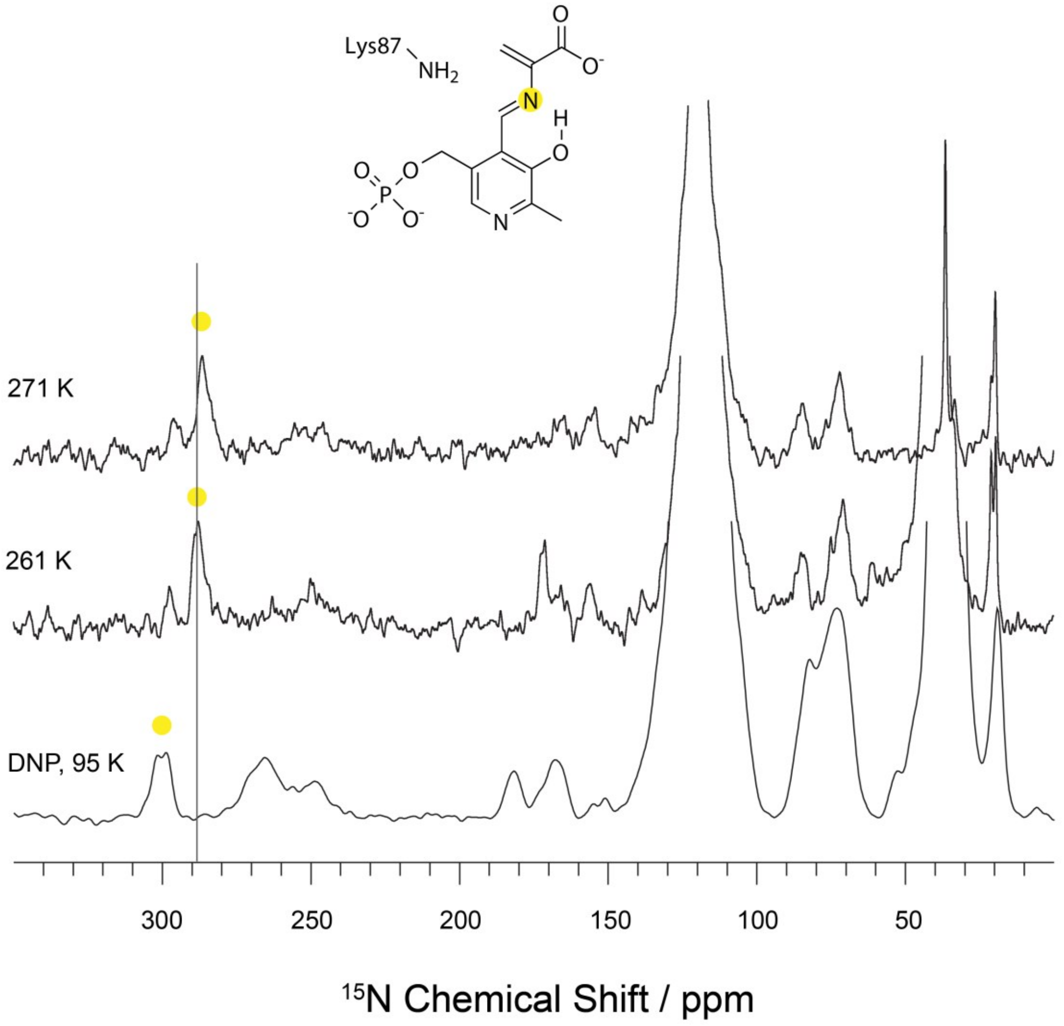


Fig. S5A. Variable-temperature $^{15}N$ SSNMR spectra of the reaction of tryptophan synthase microcrystals and $^{15}N$-Ser to form the E(A-A) intermediate. 261 K and 271 K data acquired at 14.1 T (600.01 MHz $^{1}H$, 60.82 MHz $^{15}N$) at 8 kHz MAS on a Bruker NEO spectrometer equipped with HX double resonance 4mm MAS probe. 95 K data was collected using DNP-MAS at 14.1 T on an Avance III DNP NMR system (Bruker Biospin) equipped with a Bruker 3.2 mm, $^{1}H/^{13}C/^{15}N$ MAS-DNP probe. Microwaves were delivered using a gyrotron source operating at 395 GHz and output optimized for ~12 W at the probe base. An AsymPolPOK (33, 34) concentration of ~5 mM was used, resulting in enhancements >60 and buildup times on the order of 3 s.

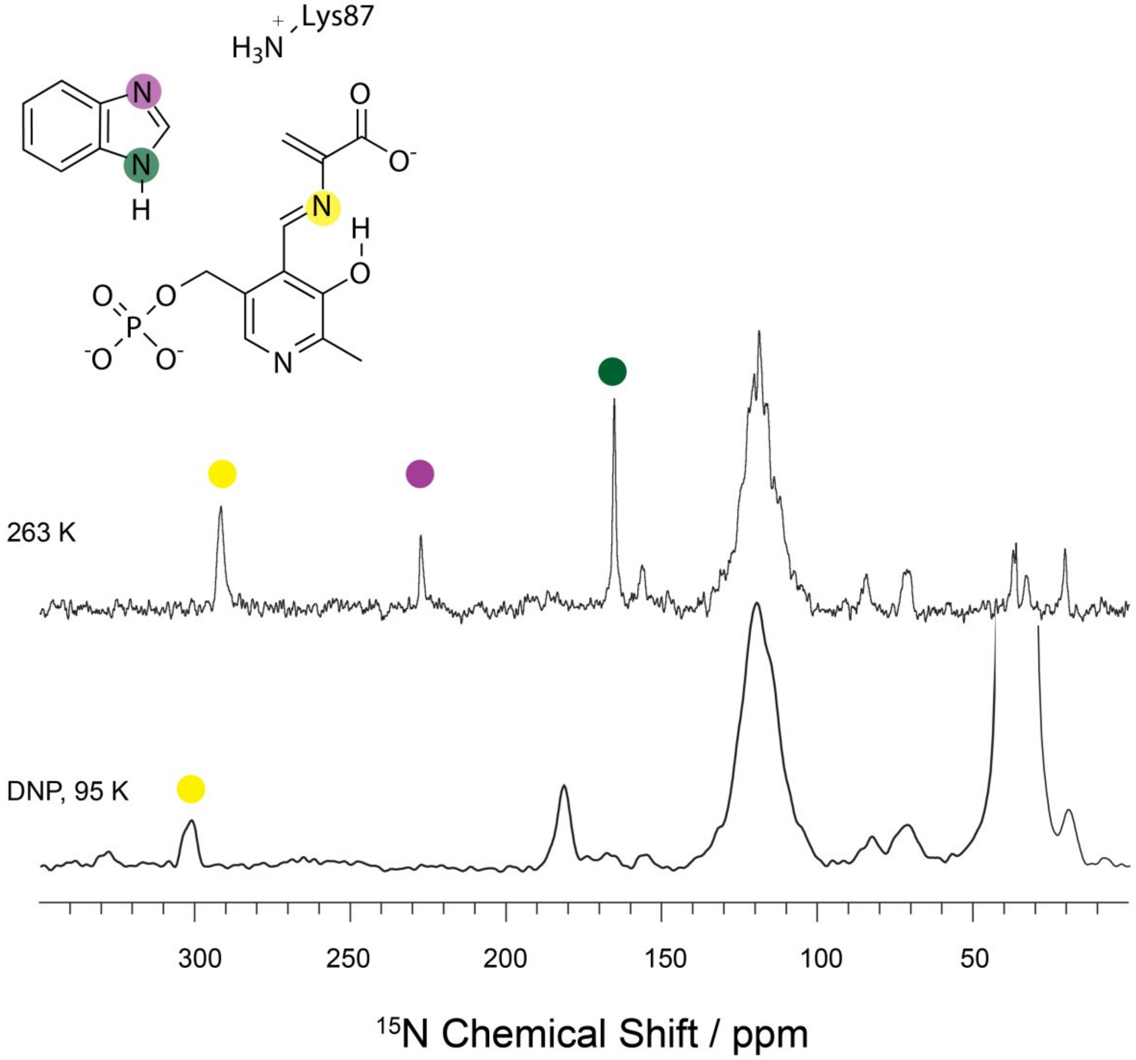


Fig. S5B. Variable-temperature $^{15}$N SSNMR spectra of the reaction of tryptophan synthase microcrystals with $^{15}$N-Ser and BZI to form the E(A-A)(BZI) complex. 263 K data acquired at 14.1 T (600.01 MHz $^{1}$H, 60.82 MHz $^{15}$N) at 8 kHz MAS on a Bruker NEO spectrometer equipped with HX double resonance 4mm MAS probe. 95 K data was collected using DNP-MAS at 14.1 T on an Avance III DNP NMR system (Bruker Biospin) equipped with a Bruker 3.2 mm, $^{1}$H/$^{13}$C/$^{15}$N MAS-DNP probe. Microwaves were delivered using a gyrotron source operating at 395 GHz and output optimized for ~12 W at the probe base. An AsymPolPOK (33, 34) concentration of ~5 mM was used, resulting in enhancements >60 and buildup times on the order of 3 s.

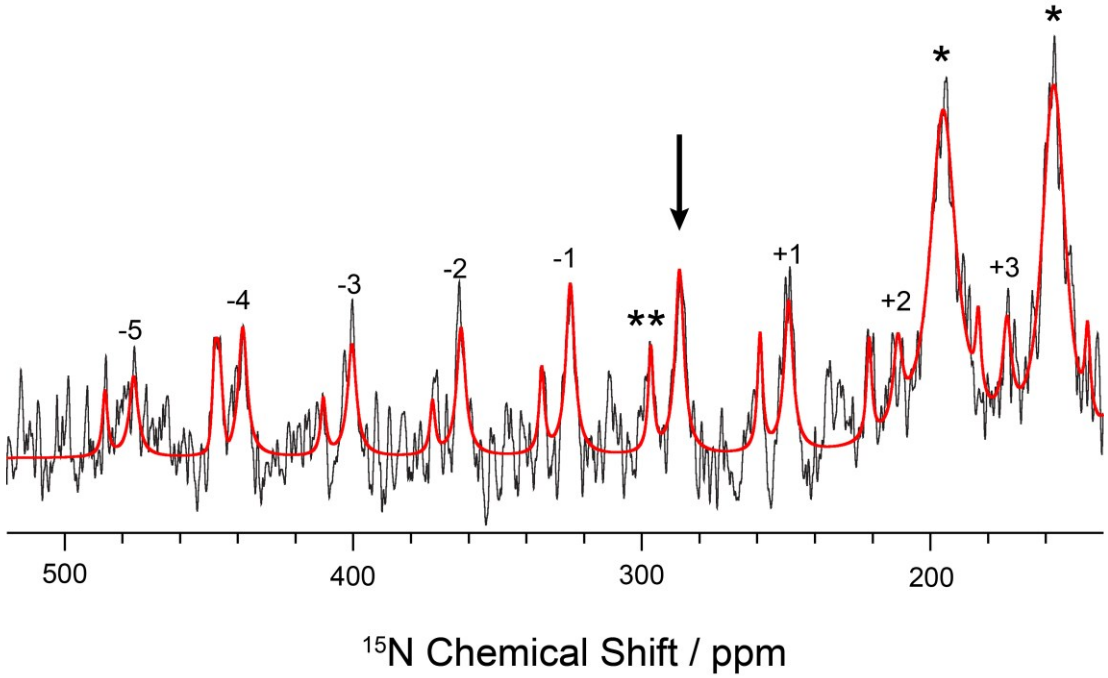


Fig. S6A. The E(A-A) Schiff base nitrogen chemical shift tensor. Slow spinning $^{15}N$ SSNMR spectrum of the reaction of tryptophan synthase microcrystals and $^{15}N$-Ser to form the E(A-A) intermediate. Experiments were performed at 14.1 T (600.01 MHz $^{1}H$, 60.82 MHz $^{15}N$) on a Bruker NEO spectrometer equipped with $^{1}H$-X double resonance 4mm MAS probe, spinning at a MAS rate of 2.3 kHz, and with the bearing gas cooled to -15°C, giving an effective sample temperature of -10°C. The primary signals arise from the isotropic shift and spinning sidebands (with sideband order given above) of the Schiff base nitrogen (286.7 ppm, indicated by arrow) and natural abundance amide backbone nitrogens (122 ppm, sidebands indicated by asterisks). A fit to the sideband manifold (red) performed in Bruker Topspin 3.6 allows for the extraction of the chemical shift principal axis components ($\delta_{11}$, $\delta_{22}$, $\delta_{33}$) = (526.5, 306.3, 31.5) ppm. There is a second minor peak present, indicated by the double asterisks.

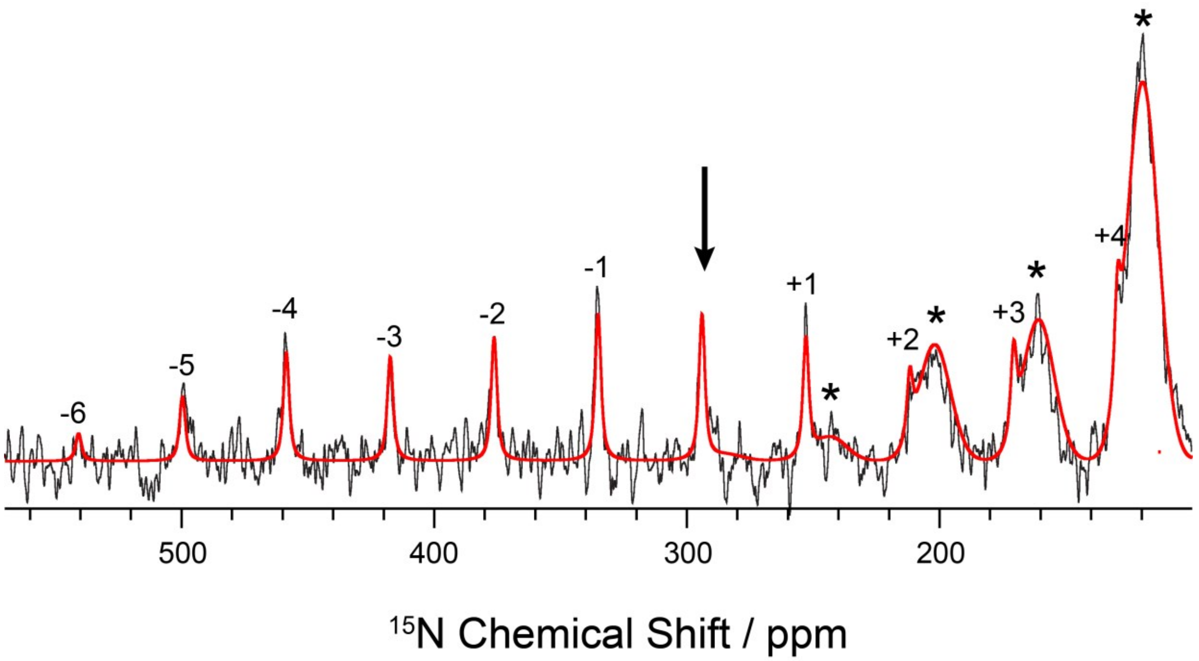


Fig. S6B. The E(A-A)(BZI) Schiff base nitrogen chemical shift tensor. Slow spinning $^{15}N$ SSNMR spectrum of the reaction of tryptophan synthase microcrystals with BZI and $^{15}N$-Ser to form the E(A-A)(BZI) complex. Experiments were performed at 14.1 T (600.01 MHz $^{1}H$, 60.82 MHz $^{15}N$) on a Bruker NEO spectrometer equipped with $^{1}H$-X double resonance 4mm MAS probe, spinning at a MAS rate of 2.5 kHz, and with the bearing gas cooled to -15°C, giving an effective sample temperature of -10°C. The primary signals arise from the isotropic shift and spinning sidebands (with sideband order given above) of the Schiff base nitrogen (292.3 ppm, indicated by arrow) and natural abundance amide backbone nitrogens (122 ppm, sidebands indicated by asterisks). A fit to the sideband manifold (red) performed in Bruker Topspin 3.6 allows for the extraction of the chemical shift principal axis components ($\delta_{11}$, $\delta_{22}$, $\delta_{33}$) = (545.8, 328.4, 7.9) ppm.

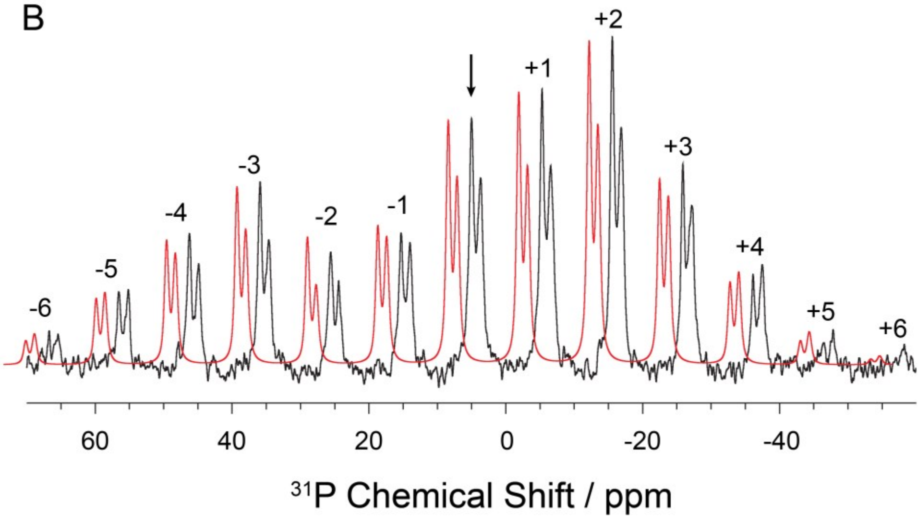


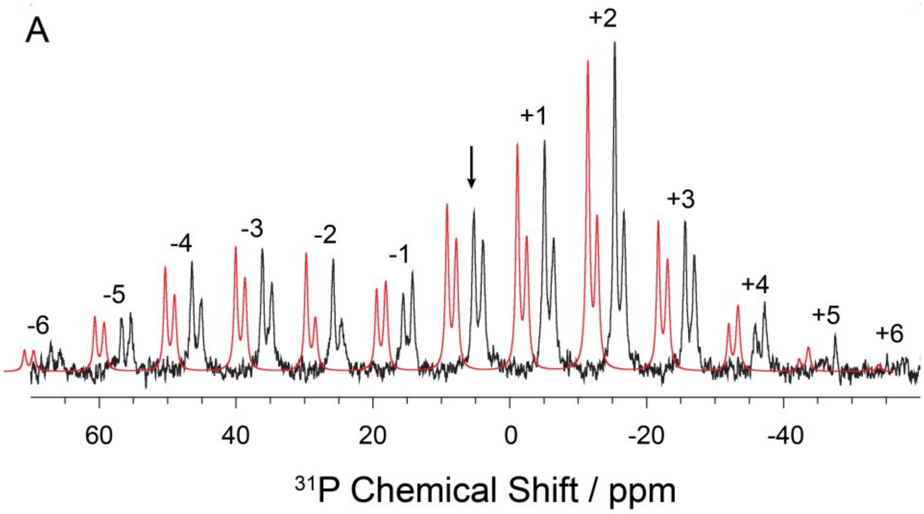


Fig. S7. The E(A-A) and E(A-A)(BZI) phosphorus chemical shift tensor. Slow spinning $^{31}P$ SSNMR CPMAS spectrum of the TS E(A-A) intermediate and E(A-A)(BZI) complex formed upon reaction TS microcrystals with L-Ser and L-Ser+BZI, respectively. The PLP phosphate isotropic peak at (A) 5.2 and (B) 4.9 ppm is indicated by the arrow. The peak at 3.7 ppm is ascribed to the α-site ligand F9. The fit (red) to the sideband manifold in BrukerTopspin 3.6 allows for the extraction of the CSA principal axis components (A) ($\delta_{11}$, $\delta_{22}$, $\delta_{33}$) = (64.6,-14.2,-34.8) ppm and (B) (A) ($\delta_{11}$, $\delta_{22}$, $\delta_{33}$) = (62.4,-8.2,-39.6) ppm for the PLP phosphate group. Both the isotropic and anisotropic chemical shifts indicate that the phosphate group is dianionic. The order of the spinning sidebands is given above each peak.

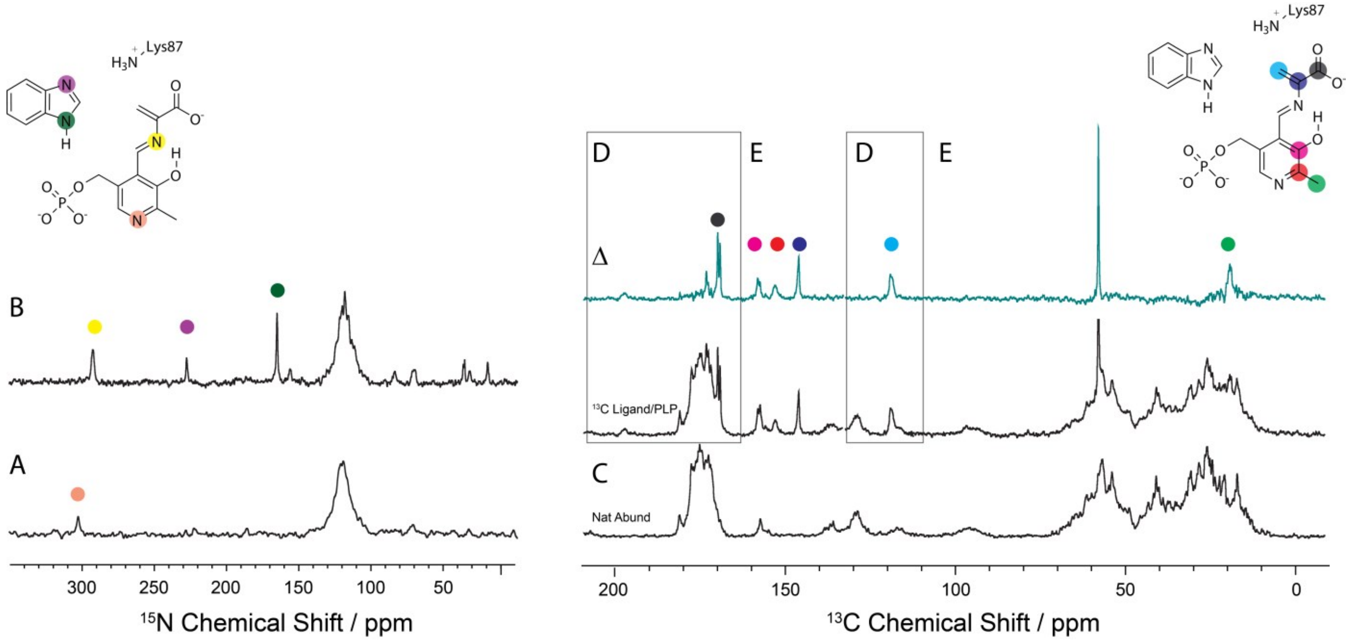


Fig. S8. $^{15}$N and $^{13}$C SSNMR CPMAS spectra of microcrystalline TS E(A-A)(BZI) prepared with the following isotopic labeling: (A) $^{15}$N-labeled on the substrate L-Ser; (B) selectively $^{15}$N enriched on the PLP cofactor and BZI; (C) natural abundance isotopomer concentration; (D) U-$^{13}C_3$-labeled L-Ser substrate; and (E) selectively $^{13}$C,$^{15}$N-enriched on the PLP cofactor and Cβ of the substrate L-Ser. The top spectra in (D)-(E) are formed as the difference between the E(A-A)(BZI) spectra with various cofactor/ligand isotopic labels and the same spectra acquired at natural abundance, highlighting the resonances for the specific site labels. The large peak at 63.1 ppm is free serine. Spectra acquired at 9.4 T, -10 °C, and 8 kHz MAS as described in the Methods and Materials.

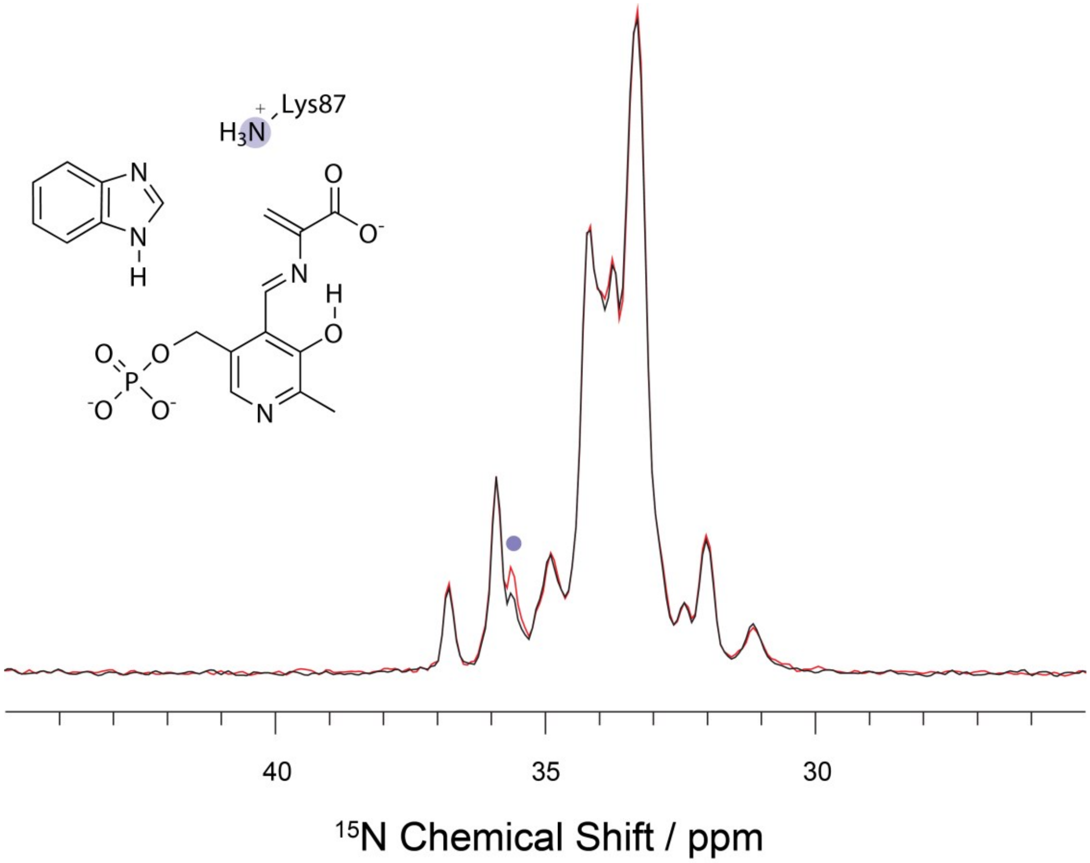


Fig. S9. $^{15}$N-observe, $^{31}$P-dephased Rotational Echo Double Resonance (REDOR) experiments of the E(A-A)(BZI) complex prepared with U-$^{15}$N-Lys TS. Black and red spectra form an $^{15}$N($^{31}$P)-REDOR pair; both have a 25 ms echo period on $^{15}$N before detection, but the black spectrum includes the application of dipolar dephasing pulses on $^{31}$P. The additional dephasing pulses specifically edit out (dephase) resonances for $^{15}$N atoms that are dipolar coupled to $^{31}$P atoms in the active site(s). The dipolar coupling falls off as the inverse cube of the interatomic distance, and the $^{15}$N($^{31}$P)-REDOR editing used here (with 25 ms of dipolar dephasing) is selective for nitrogen atoms within ~3-4 Å of the PLP $^{31}$P atoms. There is remarkable resolution of many individual ε-amino group nitrogen sites, but only the signal at 35.6 ppm dephases. Based on proximity to the PLP phosphate group, this resonance is assigned to the ε-amino group of βLys87 and, given its chemical shift, it is anticipated that it is positively charged.

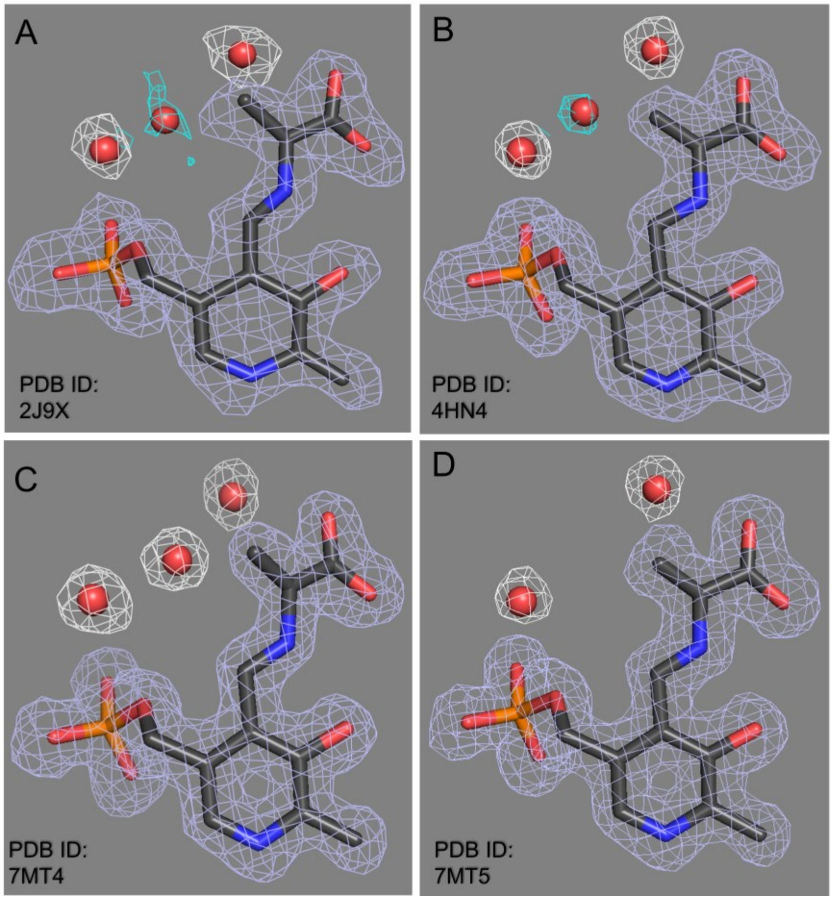


Fig. S10. Electron density maps for the E(A-A) intermediate contoured at 1 σ (white, grey) and 0.5 σ (aqua) for crystal structures (A) 2J9X, (B) 4HN4, (C) 7MT4, and (D) 7MT5.

Table S3. Statistics for X-ray crystallographic data collection and refinement

| **PDB**<br>α chain ligand<br>β chain ligands | 7MT4<br>F9F<br>$NH_4^+$, 0JO(AA) | 7MT5<br>F9F<br>$Cs^+$, 0JO(AA) | 7MT6<br>F9F<br>$Cs^+$, 0JO(AA), BZI |
|---|---|---|---|
| **Data collection** | | | |
| Detector | ADSC 315 | ADSC 315 | Rigaku RAxis V++ |
| Space group | C 1 2 1 | C 1 2 1 | C 1 2 1 |
| Unit cell, a, b, c (Å) | 183.09, 59.40, 67.34 | 184.19, 59.73, 67.254 | 183.96, 60.96, 67.41 |
| Unit cell, α, β, γ (°) | 90.00, 94.91, 90.00 | 90.00, 94.63, 90.00 | 90.00, 94.6, 90.00 |
| Wavelength (Å) | 1.00 | 1.00 | 1.54 |
| Wilson B Value ($Å^2$) | 13.37 | 12.17 | 19.8 |
| Resolution (Å) | 39.31 - 1.40<br>(1.48 -1.40) | 29.86 - 1.50<br>(1.58 - 1.50) | 29.28 - 1.70<br>(1.79 - 1.70) |
| $R_{sym}$ | 0.075 (0.331) | 0.058 (0.257) | 0.060 (0.267) |
| $R_{meas}$ | 0.082 (0.363) | 0.070 (0.308) | 0.070 (0.314) |
| $R_{pim}$ | 0.033 (0.146) | 0.038 (0.168) | 0.036 (0.164) |
| I / σI | 12.1 (4.0) | 20.9 (3.2) | 13.5 (5.1) |
| CC1/2 | 0.997 (0.950) | 0.995 (0.969) | 0.998 (0.967) |
| Completeness (%) | 95.1 (95.0) | 92.5 (86.5) | 98.5 (96.5) |
| Redundancy | 5.6 (5.6) | 2.9 (3.0) | 3.6 (3.6) |
| **Refinement** | | | |
| Resolution (Å) | 36.31-1.40 | 28.25 -1.50 | 39.23 -1.70 |
| Reflections ($R_{Free}$) | 134427 (6721) | 108105 (5240) | 80523 (4041) |
| $R_{work}$ / $R_{free}$ [a,b] (%) | 17.74 / 20.25 | 17.94 / 20.85 | 16.02 / 19.26 |
| Number of Atoms<br>(Protein, Ligand, Water) | 5777<br>(5042, 44, 691) | 5777<br>(5038, 46, 693) | 5825<br>(5052, 63, 710) |
| Average B factor ($Å^2$)<br>(Protein, Ligand, Water) | 18.95<br>(17.72, 15.67, 28.13) | 18.95<br>(17.56, 15.73, 29.24) | 23.47<br>(22.21, 21.82, 32.57) |
| R.M.S. Bond lengths (Å) | 0.007 | 0.007 | 0.007 |
| R.M.S. Bond angles (°) | 1.147 | 1.141 | 1.121 |
| Ramachandran plot (%) | | | |
| Most Favorable | 98.19 | 97.74 | 97.90 |
| Allowed | 1.66 | 2.11 | 1.95 |
| Outliers | 0.15 | 0.15 | 0.15 |

Values in parentheses are for highest-resolution shell.

[a] *R*work = Σhkl |Fo(hkl) − Fc(hkl)|/Σhkl Fo(hkl).

[b] *R*free was calculated for a test set of reflections (5%) omitted from the refinement.

### References


1. B. G. Caulkins *et al.*, NMR Crystallography of a Carbanionic Intermediate in Tryptophan Synthase: Chemical Structure, Tautomerization, and Reaction Specificity. *J Am Chem Soc* **138**, 15214-15226 (2016).
2. D. A. Case *et al.* (2018) Amber 2018. (University of California, San Francisco).
3. R. Salomon-Ferrer, A. W. Gotz, D. Poole, S. Le Grand, R. C. Walker, Routine Microsecond Molecular Dynamics Simulations with AMBER on GPUs. 2. Explicit Solvent Particle Mesh Ewald. *J Chem Theory Comput* **9**, 3878-3888 (2013).
4. D. Niks *et al.*, Allostery and substrate channeling in the tryptophan synthase bienzyme complex: evidence for two subunit conformations and four quaternary states. *Biochemistry* **52**, 6396-6411 (2013).
5. J. A. Maier *et al.*, ff14SB: Improving the Accuracy of Protein Side Chain and Backbone Parameters from ff99SB. *J Chem Theory Comput* **11**, 3696-3713 (2015).
6. J. Wang, R. M. Wolf, J. W. Caldwell, P. A. Kollman, D. A. Case, Development and testing of a general amber force field. *J Comput Chem* **25**, 1157-1174 (2004).
7. A. Jakalian, D. B. Jack, C. I. Bayly, Fast, efficient generation of high-quality atomic charges. AM1-BCC model: II. Parameterization and validation. *J Comput Chem* **23**, 1623-1641 (2002).
8. H. Nguyen, D. R. Roe, C. Simmerling, Improved Generalized Born Solvent Model Parameters for Protein Simulations. *J Chem Theory Comput* **9**, 2020-2034 (2013).
9. D. R. Roe, T. E. Cheatham, 3rd, PTRAJ and CPPTRAJ: Software for Processing and Analysis of Molecular Dynamics Trajectory Data. *J Chem Theory Comput* **9**, 3084-3095 (2013).
10. W. Humphrey, A. Dalke, K. Schulten, VMD: visual molecular dynamics. *J Mol Graph* **14**, 33-38, 27-38 (1996).
11. M. J. Frisch *et al.* (2009) Gaussian 09. (Gaussian, Inc., Wallingford, CT, USA).
12. S. Grimme, J. Antony, S. Ehrlich, H. Krieg, A consistent and accurate ab initio parametrization of density functional dispersion correction (DFT-D) for the 94 elements H-Pu. *J. Chem. Phys.* **132**, 154104 (2010).
13. J. D. Hartman, R. A. Kudla, G. M. Day, L. J. Mueller, G. J. Beran, Benchmark fragment-based (1)H, (13)C, (15)N and (17)O chemical shift predictions in molecular crystals. *Phys. Chem. Chem. Phys.* **18**, 21686-21709 (2016).
14. J. D. Hartman, T. J. Neubauer, B. G. Caulkins, L. J. Mueller, G. J. Beran, Converging nuclear magnetic shielding calculations with respect to basis and system size in protein systems. *J. Biomol. NMR* **62**, 327-340 (2015).
15. S. Moon, D. A. Case, A comparison of quantum chemical models for calculating NMR shielding parameters in peptides: mixed basis set and ONIOM methods combined with a complete basis set extrapolation. *J Comput Chem* **27**, 825-836 (2006).
16. C. W. Garland, J. W. Nibler, D. P. Shoemaker, *Experiments in physical chemistry* (McGraw-Hill Higher Education, Boston, ed. 8th, 2009), pp. x, 734 p.
17. B. G. Caulkins *et al.*, Protonation states of the tryptophan synthase internal aldimine active site from solid-state NMR spectroscopy: direct observation of the protonated Schiff base linkage to pyridoxal-5'-phosphate. *J Am Chem Soc* **136**, 12824-12827 (2014).
18. B. G. Caulkins *et al.*, Catalytic roles of betaLys87 in tryptophan synthase: (15)N solid state NMR studies. *Biochim Biophys Acta* **1854**, 1194-1199 (2015).
19. H. Ngo *et al.*, Synthesis and characterization of allosteric probes of substrate channeling in the tryptophan synthase bienzyme complex. *Biochemistry* **46**, 7713-7727 (2007).
20. E. Hilario *et al.*, Visualizing the tunnel in tryptophan synthase with crystallography: Insights into a selective filter for accommodating indole and rejecting water. *Biochim Biophys Acta* **1864**, 268-279 (2016).
21. M. D. Winn *et al.*, Overview of the CCP4 suite and current developments. *Acta Crystallogr D Biol Crystallogr* **67**, 235-242 (2011).
22. P. Emsley, K. Cowtan, Coot: model-building tools for molecular graphics. *Acta Crystallogr D Biol Crystallogr* **60**, 2126-2132 (2004).
23. G. N. Murshudov *et al.*, REFMAC5 for the refinement of macromolecular crystal structures. *Acta Crystallogr D Biol Crystallogr* **67**, 355-367 (2011).
24. D. Liebschner *et al.*, Macromolecular structure determination using X-rays, neutrons and electrons: recent developments in Phenix. *Acta Crystallogr D Struct Biol* **75**, 861-877 (2019).

25. A. Peracchi, A. Mozzarelli, G. L. Rossi, Monovalent cations affect dynamic and functional properties of the tryptophan synthase. alpha. 2. beta. 2 complex. *Biochemistry* **34**, 9459-9465 (1995).
26. E. Woehl, M. F. Dunn, Mechanisms of monovalent cation action in enzyme catalysis: the tryptophan synthase α-, β-, and αβ-reactions. *Biochemistry* **38**, 7131-7141 (1999).
27. A. T. Dierkers, D. Niks, I. Schlichting, M. F. Dunn, Tryptophan Synthase: Structure and Function of the Monovalent Cation Site. *Biochemistry* **48**, 10997-11010 (2009).
28. B. M. Fung, A. K. Khitrin, K. Ermolaev, An improved broadband decoupling sequence for liquid crystals and solids. *J. Magn. Reson.* **142**, 97-101 (2000).
29. R. K. Harris *et al.*, Further conventions for NMR shielding and chemical shifts (IUPAC Recommendations 2008). *Pure and Applied Chemistry* **80**, 59-84 (2008).
30. C. R. Morcombe, K. W. Zilm, Chemical shift referencing in MAS solid state NMR. *J. Magn. Reson.* **162**, 479-486 (2003).
31. T. Maurer, H. R. Kalbitzer, Indirect Referencing of 31P and 19F NMR Spectra. *J Magn Reson B* **113**, 177-178 (1996).
32. T. Gullion, J. Schaefer, Rotational-Echo Double-Resonance Nmr. *Journal of Magnetic Resonance* **81**, 196-200 (1989).
33. S. A. McNeill, P. L. Gor'kov, K. Shetty, W. W. Brey, J. R. Long, A low-E magic angle spinning probe for biological solid state NMR at 750 MHz. *J. Magn. Reson.* **197**, 135-144 (2009).
34. B. M. Fung, A. K. Khitrin, K. Ermolaev, An improved broadband decoupling sequence for liquid crystals and solids. *J. Magn. Reson.* **142**, 97-101 (2000).
35. J. Herzfeld, A. E. Berger, Sideband Intensities in Nmr-Spectra of Samples Spinning at the Magic Angle. *Journal of Chemical Physics* **73**, 6021-6030 (1980).
36. G. Hou, S. Yan, J. Trebosc, J. P. Amoureux, T. Polenova, Broadband homonuclear correlation spectroscopy driven by combined R2(n)(v) sequences under fast magic angle spinning for NMR structural analysis of organic and biological solids. *J. Magn. Reson.* **232**, 18-30 (2013).
37. A. Hassan *et al.*, Sensitivity boosts by the CPMAS CryoProbe for challenging biological assemblies. *J. Magn. Reson.* **311**, 106680 (2020).
38. N. T. Tran, F. Mentink-Vigier, J. R. Long, Dynamic Nuclear Polarization of Biomembrane Assemblies. *Biomolecules* **10**, 1246 (2020).
39. E. S. Salnikov *et al.*, Dynamic Nuclear Polarization/Solid-State NMR Spectroscopy of Membrane Polypeptides: Free-Radical Optimization for Matrix-Free Lipid Bilayer Samples. *Chemphyschem* **18**, 2103-2113 (2017).
40. X. Wang *et al.*, Direct dynamic nuclear polarization of 15N and 13C spins at 14.1 T using a trityl radical and magic angle spinning. *Solid state nuclear magnetic resonance* **100**, 85-91 (2019).
41. Z. Gan *et al.*, NMR spectroscopy up to 35.2T using a series-connected hybrid magnet. *J. Magn. Reson.* **284**, 125-136 (2017).
42. R. P. Young *et al.*, Solution-State 17O Quadrupole Central-Transition NMR Spectroscopy in the Active Site of Tryptophan Synthase. *Angewandte Chemie International Edition* **55**, 1350-1354 (2016).
43. P. P. Man, J. Klinowski, A. Trokiner, H. Zanni, P. Papon, Selective and Non-Selective Nmr Excitation of Quadrupolar Nuclei in the Solid-State. *Chemical Physics Letters* **151**, 143-150 (1988).
44. J. F. Zhu, G. Wu, Quadrupole Central Transition O-17 NMR Spectroscopy of Biological Macromolecules in Aqueous Solution. *Journal of the American Chemical Society* **133**, 920-932 (2011).
45. C. V. Grant *et al.*, A Modified Alderman-Grant Coil makes possible an efficient cross-coil probe for high field solid-state NMR of lossy biological samples. *Journal of Magnetic Resonance* **201**, 87-92 (2009).
46. R. Fu *et al.*, Ultra-wide bore 900MHz high-resolution NMR at the National High Magnetic Field Laboratory. *Journal of Magnetic Resonance* **177**, 1-8 (2005).
47. M. S. Ghatge *et al.*, Pyridoxal 5'-phosphate is a slow tight binding inhibitor of E. coli pyridoxal kinase. *PLoS One* **7**, e41680 (2012).
48. D. F. Gauto *et al.*, Integrated NMR and cryo-EM atomic-resolution structure determination of a half-megadalton enzyme complex. *Nat Commun* **10**, 2697 (2019).
49. J. Shen *et al.*, A Quadrupole-Central-Transition 17O NMR Study of Nicotinamide: Experimental Evidence of Cross-Correlation between Second-Order Quadrupolar Interaction and Magnetic Shielding Anisotropy. *The Journal of*

*Physical Chemistry B* **122**, 4813-4820 (2018).
50. G. Wu, 17O NMR studies of organic and biological molecules in aqueous solution and in the solid state. *Progress in Nuclear Magnetic Resonance Spectroscopy* **114-115**, 135-191 (2019).
51. L. G. Werbelow, Nmr Dynamic Frequency-Shifts and the Quadrupolar Interaction. *Journal of Chemical Physics* **70**, 5381-5383 (1979).
